# Travelling Waves in Gene Expression: A Mathematical Model of Cell-State Dynamics in Melanoma

**DOI:** 10.1101/2025.10.18.683212

**Authors:** Charlotte Taylor Barca, Rotem Leshem, Vishaka Gopalan, Sarah Woolner, Kerrie L. Marie, Gareth Wyn Jones, Oliver E. Jensen

## Abstract

Melanoma is a cancer of the melanocyte, known to have an ability to readily switch between different transcriptional cell states that convey different phenotypic properties (e.g. hyper-differentiated, neural crest-like). This ability is believed to underpin intratumour heterogeneity and plastic adaptation, which contributes to resistance to therapy and immune evasion of the tumour. Therefore, understanding the mechanisms underlying acquisition of transcriptional cell states and cell-state switching is crucial for the development of therapies. We model a minimal gene regulatory network comprising three key transcription factors, whose varying gene expression encodes different melanoma cell states, and use deterministic spatiotemporal differential-equation models to study gene-expression dynamics. We exploit an approximation, based on cooperative binding of transcription factors, in which the models are piecewise-linear. We classify stable states of the local model in a biologically relevant manner and, using a naïve model of intercellular communication, we explore how a population of cells can take on a shared characteristic through travelling waves of gene expression. We derive a condition determining which characteristic will become dominant, under sufficiently strong cell-cell signalling, which creates a partition of parameter space.

## 1 Introduction

A tumour’s ability to plastically adapt (through cell-state switching) to new environments underpins its ability to both metastasise and resist therapeutic intervention. Heterogeneous tumours consist of cells belonging to many different transcriptional cell states, which can encode different cellular phenotypes. The heterogeneity of melanoma (a cancer of pigment-producing melanocytes in the skin) has been linked to its ability to resist therapies and evade the immune system [1]. The complexity of a heterogeneous tumour and the dynamic nature of cells capable of state-switching make it difficult to map and predict the evolution of tumour cells during melanoma progression. The main method for studying cell states in melanoma tumours and cells is the measurement of protein or RNA levels of key cell-state markers by taking snapshots of relative levels at a given time, and building understanding of the underlying regulatory networks. Central to these regulatory processes are transcription factors (TFs), proteins that attach to promoter or enhancer sequences in DNA and activate or repress the expression of genes associated with that promoter. Gene regulatory networks (GRNs) can involve hundreds of genes and TFs, making the analysis of such systems a challenging task. Here, we study a refined version of a GRN in melanoma and use mathematical modelling to explore cell-state switches within a cell, and, how groups of interacting cells may influence each other’s states. Adopting a dynamical-systems perspective, we analyse a non-smooth approximation of the system’s governing equations and identify a finite set of steady states. Our analysis reveals bistable regions of parameter space, suggests potential state-transition pathways via saddle-node bifurcations, and highlights key bifurcation parameters.

The concept of phenotypic state switching in melanoma has been strongly influenced through the work of Goding [2–4], who discovered that variable levels of the melanocyte transcription factor, MITF, controlled a switch between proliferative and invasive phenotypes in melanoma [2]. Later, greater granularity was brought to this concept, ultimately suggesting that high levels of MITF induced hyper-differentiated, quiescent-like states that can be tolerant to therapy, low levels of MITF induced invasion, and medium levels drove proliferation [3–5]. Beyond MITF, we now understand that it is the multiple and interacting TF networks that influence the phenotypes in melanoma [2, 6–8]. A GRN that has been implicated is a MITF, SOX10 and ZEB1 network [9–15]. SOX10 is a TF that specifies melanocyte precursors from the embryonic neural crest. High levels of SOX10 (along with medium levels of MITF) can drive a proliferative phenotype. ZEB1 is a TF that drives an epithelial–mesenchymal transition; it can directly inhibit MITF and SOX10 expression and is known to promote an invasive mesenchymal-like cell state or a neural stem cell-like state [9, 15]. MITF has been shown in turn to negatively regulate ZEB1, impacting cell morphology and cell-to-environment interactions [14]. While this is a simplification of what is known to be a complex GRN, we can use these phenotypic classifications to identify emergent cell states in a mathematical model.

Beyond this local understanding of gene expression, we are also motivated by tumour-level observations of regions of cell-state clustering in melanoma, potentially implicating intercellular communication. Multiple studies have reported that cells in specific regions of a tumour can exhibit similar expression markers [6, 16–18], yet across the tumour there is heterogeneity between different regions. For example, one metastatic lesion may show high SOX10 expression and low ZEB1 expression, while a nearby metastatic lesion may display the reverse pattern, with other areas showing mixed markers. Figure 1 shows immuno-stained images of subcutaneous melanoma tumours, in which cells have been stained for markers of each TF or TF pathway activation. All SOX10 positive cells in Figure 1 are tumour cells. Cells in the central oval-shaped region of the tumour have high SOX10, low ZEB1 and high MITF pathway activation, whereas the remaining tumour cells exhibit the opposite behaviour, raising the question of which mechanisms drive the heterogeneity observed across the tumour.

**Fig. 1.**
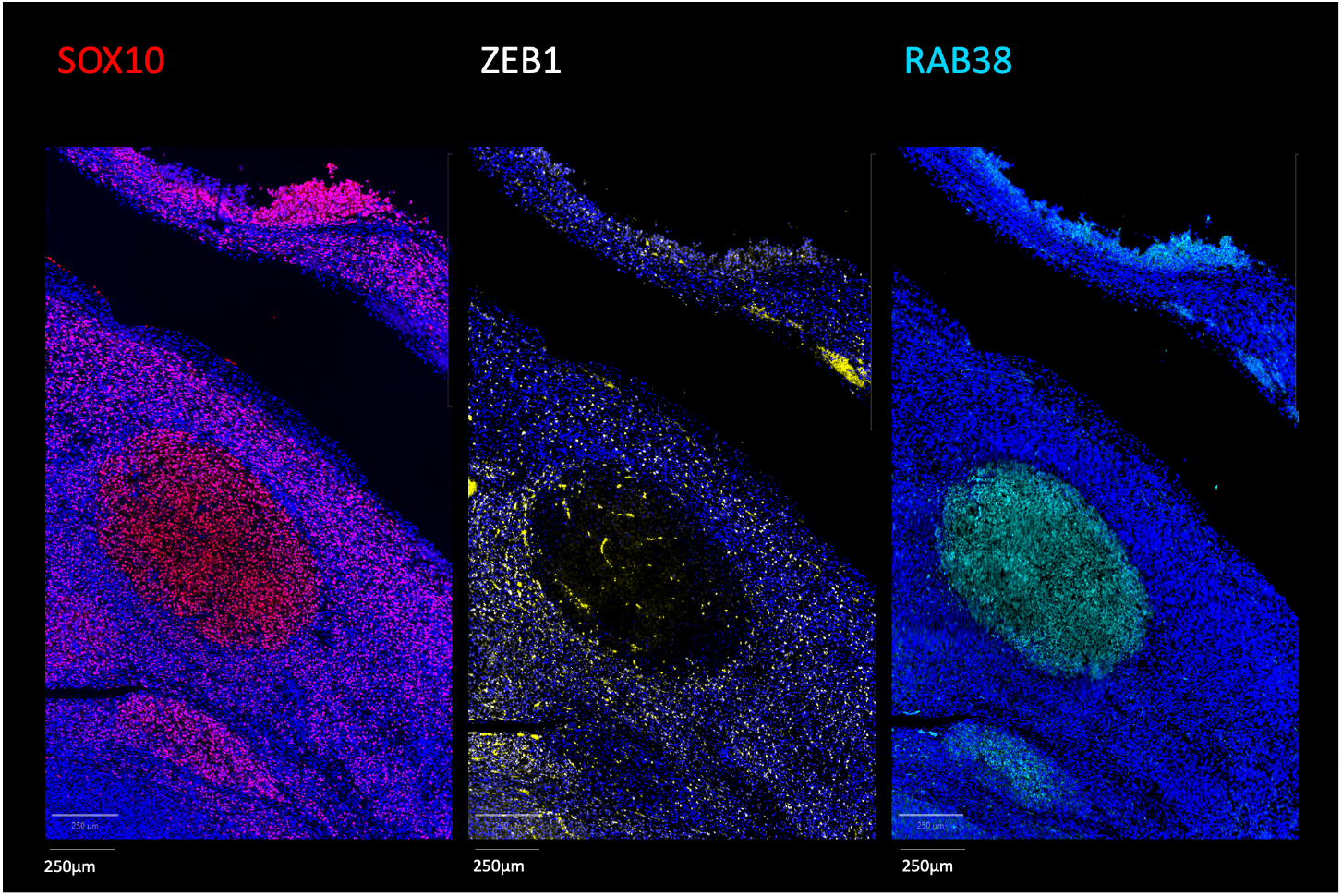
M4 mouse melanoma tumour experimentally induced by transplantation of B2905 melanoma cells subcutaneously into immune-competent mice. Images are taken from a single plane. Immunostaining shows SOX10 (left), ZEB1 (middle) and RAB38 expression, a gene involved in melanosome biogenesis and maturation, whose expression is directly activated by MITF binding, and which therefore acts as a readout for MITF activity (right). The true ZEB1 TF staining appears white in the nuclei, whereas the pure yellow is a result of a blood vessel artefact. All cells displaying SOX10 expression are tumour cells. Dark blue staining across the three images is DAPI, a nuclear marker. Images taken by Rotem Leshem.

We base our modelling approach on a model proposed by Subhadarshini *et al*. [19], who described a GRN in melanoma comprising five TFs (SOX10, MITF, SOX9, JUN, ZEB1) using coupled ordinary differential equations (ODEs). Subhadarshini *et al*. identified stable steady states by solving the ODE system over randomly generated initial conditions and parameter values (sampled from biologically feasible ranges). By hierarchical clustering, they identified four distinct phenotypic cell states: undifferentiated, neural-crest like, intermediate and proliferative. Subhadarshini *et al*. used these observations, alongside transcriptomic data analysis, to investigate the impact of state heterogeneity in therapy evasion.

We build on [19] by (i) adding an inhibitory link from ZEB1 to SOX10, based on recent advances in the field [9, 10]; (ii) reducing the GRN to

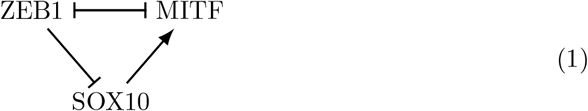

by discounting SOX9 and JUN from the network; (iii) reducing the model to piecewise-linear form under the assumption that large numbers of TFs bind to promoter sites; and (iv) incorporating spatial structure. Piecewise-linear models can be used to model GRNs [20, 21] by simplifying the commonly-used Hill function [22], allowing consideration of the stability of states by partitioning the phase plane [20]. We exclude SOX9 and JUN from the network in [19] because their regulatory roles are restricted to interactions with SOX10 and MITF, respectively. Their effects can be considered secondary and are absorbed into the dynamics of SOX10 and MITF. To account for spatial effects, we model a multicellular tissue as a Voronoi tessellation, capturing the disordered nature of cell-packing (as seen in Figure 1). We model intercellular communication via diffusion. Concentration gradients of diffusible signalling molecules across a population of cells can induce changes in gene expression [23, 24]. We model the resulting spatial changes in gene expression (together with local GRN dynamics) using reaction-diffusion equations [25, 26]. Under this framework, we analyse travelling-wave solutions in gene expression across the cell population [27]. Such waves represent a spatial coordination of a phenotypic cell state as a response to intercellular signalling [28, 29].

Motivated by the model and existing literature, we will consider the following cell states (and the phenotypes they are associated with): hyper-differentiated (quiescent, drug-tolerant); transitory (hybrid); melanocytic (proliferating); neural-crest-like (drug tolerant, slow cycling, stem-cell like); mesenchymal-like (undifferentiated, invasive); MZ-hybrid (MITF/ZEB1 double positive); alternate; MITF^+^; and SOX10^+^. The SOX10, MITF and ZEB1 expression profiles which characterise each of these cell states in our model can be found below.

The paper is structured as follows. In Section 2 we define the model and methods we use throughout, and in Section 3 we present results. In Section 2, by analysis of a simplified system of ODEs, we study how stable states of the GRN interact with each other. To understand how cells in a lesion may influence each other, in Section 2.1 we introduce a spatial element to the model by allowing diffusion of TFs between neighbouring cells. (In our naïve model of intercellular communication, the diffusion terms represent the effect of some diffusing signalling molecules, which we do not explicitly model, to which TF levels are strongly coupled.) We study parameter regions having coexisting stable and unstable states, and identify bifurcation parameters that potentially drive state transitions in Section 2.2. Using tools from discrete calculus, we use this model to simulate interacting subpopulations of cells in different locally-uniform states in Section 3, and use a travelling-wave analysis to predict which states may dominate in a population of cells. We assess the model’s robustness to the assumption of large numbers of TFs for cooperative binding, and discover a butterfly catastrophe.

## 2 Model and Methods

We adopt a continuous-time discrete-space framework, by formulating a local ODE model derived from a reaction-based representation of TF interactions, which we extend to account for spatial structure in Section 2.1 below. In Section 2.2 we determine the steady uniform states of the system, and determine their stability to spatially-uniform perturbations. We explore parameter space to determine regions of bistability, and we set up the simulations that we consider in Section 2.3. In Section 2.4 we study interacting subpopulations of cells at steady locally-uniform states through a travelling-wave analysis.

We consider the GRN (1), a directed network of activating and inhibiting relationships between TFs in the nucleus of a melanoma cell. Denoting the TFs SOX10, MITF and ZEB1 by *S, M* and *Z*, respectively, we model the interactions

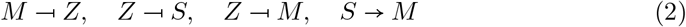

using mass-action kinetics and a simplified biological mechanism of activation and inhibition in the system. In Appendix A.1, we derive the local ODE model of dynamics of the GRN (2), from a reaction-based representation of TF interactions. We denote by 𝒮, ℳ, *Ƶ* the genes that encode TFs *S, M, Z*, respectively. We use the symbol ℬ to refer to an arbitrary gene and *α* to refer to an arbitrary TF. Each gene has associated promoter regions to which appropriate TFs can bind, impacting the production rate of the TF that the gene encodes. Activators help the RNA polymerases (enzymes which catalyse the transcription process) bind to the promoter region, whereas inhibitors may block the RNA polymerase from binding. In reality, a gene is regulated by various TFs and may have multiple promoter or enhancer regions to which these will bind. In this refined GRN framework, we simplify the regulatory mechanism by assigning to each gene a distinct promoter region for each TF that regulates it, as specified by the network (2). Other regulatory effects are absorbed into the basal production rates of each TF.

Let 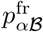 be a free space on the promoter region of gene ℬ to which TF *α* will bind, for (*α*, ℬ) ∈ {(*M, Ƶ*), (*Z*, S), (*Z*, ℳ), (*S*, ℳ)}. For gene ℬ, we refer to this as ℬ’s ‘*α*-promoter region’. When 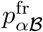 meets *n*_*α*_ copies of TF *α*, the promoter space is labelled as bound, 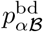, and is no longer available for binding. The TFs bind cooperatively in this sense. For simplicity we assume that *n*_*α*_ – the number of copies of TF *α* required to fully bind to the promoter region of a gene for inhibition or activation – is not dependent on the target gene, taking *n*_*S*_ = *n*_*M*_ = *n*_*Z*_ = *n* for some integer *n*. Finally, we denote by *R* the RNA polymerase that binds to a gene to initiate transcription.

We introduce parameters *λ*_*αβ*_, the fold changes in production of TF *β* as a result of activation or inhibition by TF *α*, which satisfy *λ*_*αβ*_ > 1 or *λ*_*αβ*_ < 1, respectively. We use the notation *S* = [*S*], *M* = [*M*], *Z* = [*Z*] – concentrations of *S, M* and *Z* in a cell – and the Hill-type sigmoidal function

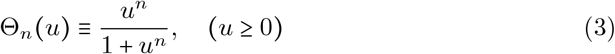

to model TF interactions, which we derive in Appendix A.1. We make use of the fact that Θ_*n*_ (*u*) increases from 0 to 1 as *u* increases from *u* ≪ 1 to *u* ≫ 1, and that Θ _−*n*_ decreases from 1 to 0. The resulting system of ODEs is

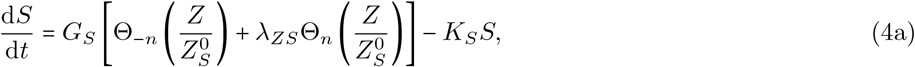

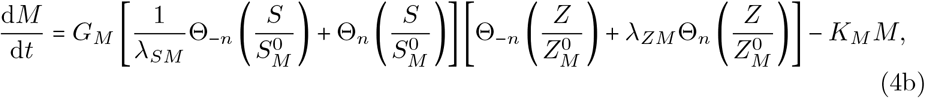

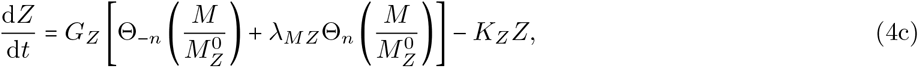

introducing production rates *G*_*S*_, *G*_*M*_, *G*_*Z*_, degradation rates *K*_*S*_, *K*_*M*_, *K*_*Z*_, thresholds 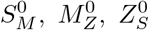 and 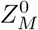 and amplitudes *λ*_*ZS*_, *λ*_*SM*_, *λ*_*ZM*_ and *λ*_*MZ*_. (Subscripts on thresholds, and the second subscript on amplitudes, denote the target of activation or inhibition.) The terms Θ_−*n*_ (⋅) and Θ_*n*_ (⋅) in (4) represent the effects on a TF’s production rate of unbound and bound promoter spaces, respectively. As *n* → ∞,

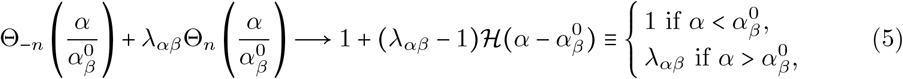

where ℋ (⋅) is the Heaviside function. In this limit the production rates of TFs are piecewise-constant, which we will exploit to analyse the system.

### 2.1 Tissue-level model

We use a Voronoi tessellation (with periodic boundary conditions) to mimic the cellular structure of a tissue. We adopt the notation in [30] (see Appendix A.2 for details) and define a discrete Laplacian ℒ (A8), for scalar fields defined over cell centres. With this and the local ODE model (4) we define the reaction-diffusion system

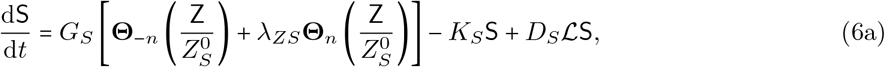

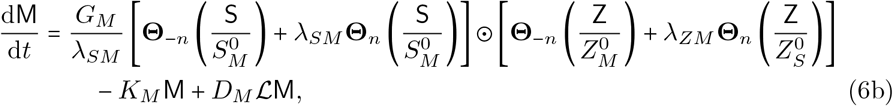

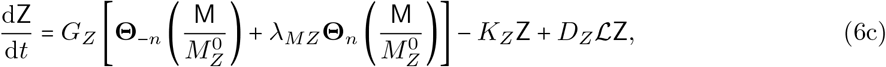

with initial conditions S 0 = S_0_, M (0) = M_0_, Z (0) = Z_0_. *D*_*S*_, *D*_*M*_, *D*_*Z*_ are diffusion coefficients; we expect diffusion to be mediated by secondary signalling molecules, which in turn impact TF expression and activity. We do not simulate these signalling molecules directly. S, M, Z and **Θ**_*n*_ *α* are vectors containing *S, M, Z* and Θ_*n*_ *α* values in each cell, respectively and ⊙ is the Hadamard (element-wise) product. Thus (6) forms 3*N*_*c*_ coupled equations with *N*_*c*_ the number of cells. The operator ℒ is positive semi-definite and satisfies ℒ1 = 0, where 1 = (1, …, 1)^⊤^, so the system (6) conserves mass. In Appendix A.2 we nondimensionalise (6) to obtain

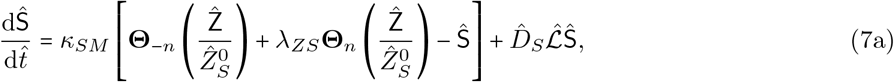

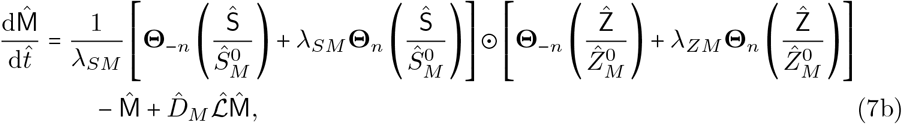

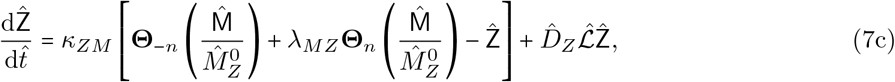

where *κ*_*SM*_ ≡ *K*_*S*_/*K*_*M*_ and *κ*_*ZM*_ ≡ *K*_*z*_*/K*_*M*_, with initial conditions Ŝ (0) Ŝ_0_, 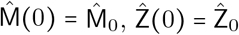. System (7) has 14 dimensionless parameters, namely *κ*_*SM*_, *κ*_*ZM*_, *λ*_*ZS*_, *λ*_*SM*_, *λ*_*ZM*_, *λ*_*MZ*_, 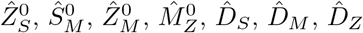 and *n*.

### 2.2 Steady uniform states

In the limit *n* → ∞, (7) can be written using (5) in the form

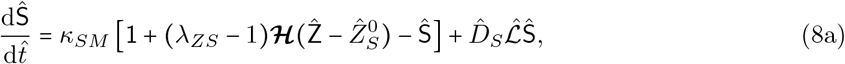

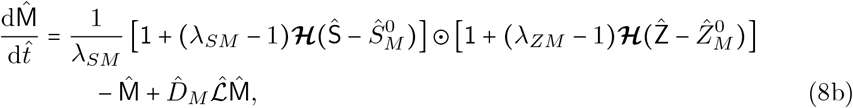

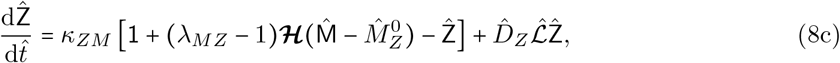

where ℋ (⋅) is the vector of Heaviside functions in each cell. We use this limit to find approximate steady uniform states of (7). With spatial homogeneity the diffusion terms vanish 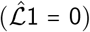, so steady uniform states are the solutions of the non-smooth algebraic system

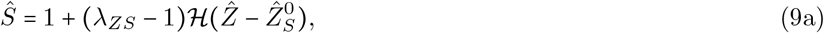

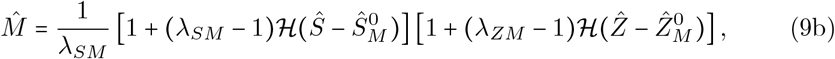

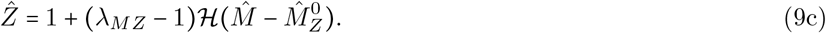

We assume throughout that *λ*_*ZM*_ *<* 1/*λ*_*SM*_ for illustration purposes. The following inequalities are consequently fixed:

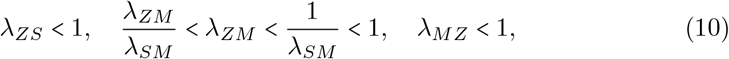

with which we categorise stable steady-state values as high/medium/low *Ŝ*, 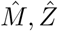 in Table 1. We formalise the description of phenotypic cell states in Section 1 as combinations of these *Ŝ*, 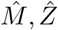 state labels in Table 2. The steady uniform states satisfying (9) (evaluated in Appendix A.3) are listed in Table 3, and the parameter restrictions on each state are listed in Tables A1–A3. The four Heaviside functions in (9) generate 2^4^ possible states; three additional states emerge at boundaries between parameter regions. The stability analysis in Appendix A.4 shows that, in the limit *n* → ∞, states 1–16 in Table 3 are stable to spatially-uniform perturbations, and states 17–19 are saddle points. Motivated by this, in Table 3 the stable states are assigned a label, grouping states into transcriptomic cell-state subtypes (and associated phenotypes) defined by criteria specified in Tables 1 and 2.

**Table 1.** Classification of steady-state values of *Ŝ*, 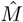 and 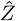 into L, M, and H categories (low, medium and high-expression states, respectively, of *Ŝ*, 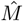 and 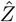), according to the restrictions (10).

| TF | Value | L/M/H |
| --- | --- | --- |
| $\hat{S}$ | $\lambda_{ZS}$ | L |
|  | 1 | H |
| $\hat{M}$ | $\lambda_{ZM}/\lambda_{SM}$ | L |
| | $\lambda_{ZM}$ | M |
| | $1/\lambda_{SM}$ | M |
|  | 1 | H |
| $\hat{Z}$ | $\lambda_{MZ}$ | L |
|  | 1 | H |

**Table 2.** Definition of cell states of interest. SOX10, ZEB1 and MITF expression levels extracted from Tsoi *et al*. [7] were used to define labels. These labels are used in Table 3 to classify the uniform steady states of the model according to their values in Table 1.

| State | SOX10 | MITF | ZEB1 |
| --- | --- | --- | --- |
| Hyper-differentiated, **HD** | L/H | H | L |
| Transitory, **TR** | H | H/M | H |
| Melanocytic, **ML** | H | M | L |
| Neural crest-like, **NC** | H | L | H |
| Mesenchymal-like, **MS** | L | L | H |
| MZ-hybrid, **HY** | L | M/H | H |
| Alternate, **AL** | L | L | L |
| MITF<sup>+</sup>, M+ | L | M | L |
| SOX10<sup>+</sup>, S+ | H | L | L |

**Table 3.** List of steady uniform states of (7), grouped by phenotype. States are categoriesed as HD, TR, ML/NC, MS, HY, AL, M+, S+, as defined in Table 2; SA denotes saddle points.

| Equilibrium Point<br>($\hat{S}, \hat{M}, \hat{Z}$) | Label |
| --- | --- |
| 1. $(\lambda_{ZS}, 1, \lambda_{MZ})$ | **HD1** |
| 2. $(1, 1, \lambda_{MZ})$ | **HD2** |
| 3. $(1, 1, 1)$ | **TR1** |
| 4. $(1, \frac{1}{\lambda_{SM}}, 1)$ | **TR2** |
| 5. $(1, \lambda_{ZM}, 1)$ | **TR3** |
| 6. $(1, \frac{1}{\lambda_{SM}}, \lambda_{MZ})$ | **ML1** |
| 7. $(1, \lambda_{ZM}, \lambda_{MZ})$ | **ML2** |
| 8. $(1, \frac{\lambda_{ZM}}{\lambda_{SM}}, 1)$ | **NC** |
| 9. $(\lambda_{ZS}, \frac{\lambda_{ZM}}{\lambda_{SM}}, 1)$ | **MS** |
| 10. $(\lambda_{ZS}, 1, 1)$ | **HY1** |
| 11. $(\lambda_{ZS}, \lambda_{ZM}, 1)$ | **HY2** |
| 12. $(\lambda_{ZS}, \frac{1}{\lambda_{SM}}, 1)$ | **HY3** |
| 13. $(\lambda_{ZS}, \frac{\lambda_{ZM}}{\lambda_{SM}}, \lambda_{MZ})$ | **AL** |
| 14. $(\lambda_{ZS}, \frac{1}{\lambda_{SM}}, \lambda_{MZ})$ | **M+1** |
| 15. $(\lambda_{ZS}, \lambda_{ZM}, \lambda_{MZ})$ | **M+2** |
| 16. $(1, \frac{\lambda_{ZM}}{\lambda_{SM}}, \lambda_{MZ})$ | **S+** |
| 17. $(\hat{S}_M^0, \hat{M}_Z^0, \hat{Z}_S^0)$ | **SA1** |
| 18. $(\lambda_{ZS}, \hat{M}_Z^0, \hat{Z}_M^0)$ | **SA2** |
| 19. $(1, \hat{M}_Z^0, \hat{Z}_M^0)$ | **SA3** |

#### 2.2.1 Bistable parameter regions

We now consider how the states in Table 3 populate parameter space. The parameter restrictions associated with each state do not form a partition of parameter space. For example, 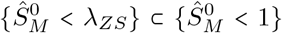. Motivated by the parameter restrictions on each state we consider where 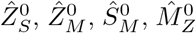 fall relative to the parameter intervals (10) to identify bistable parameter regions, allowing us to consider a four-dimensional parameter space (rather than a 13-dimensional one). We do so using the projection illustrated in Figure 2. We have chosen to project onto the subspaces 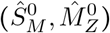 and 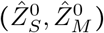 because saddle points are restricted by 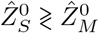 (see Tables A2, A3). The two-dimensional (2D) projection in Figure 2(a) lists the states in each region of the 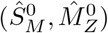-plane which satisfy restrictions on 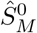 and 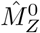 only (discounting restrictions on 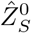 and 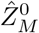). Figure 2(b) shows the states in each region of the 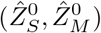-plane satisfying restrictions on 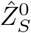 and 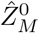 only (discounting restrictions on 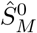 and 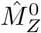). In this sense, a region of the projected parameter space is defined by pairing regions from Figures 2(a) and 2(b), as described by the steps on the Figure. Feasible steady states are found by comparing the states in each box: the intersection of these states identify temporally-stable steady state(s) for the region. For any given parameter region Ω the system exhibits either a unique temporally-stable steady state (monostable) or a pair of temporally-stable steady states (bistable) with a third saddle point. All parameters on the axes in Figure 2 can vary, but are restricted to the order in which they appear as a result of (10). Further analysis of stable states is presented in Section 3.1 below.

**Fig. 2.**
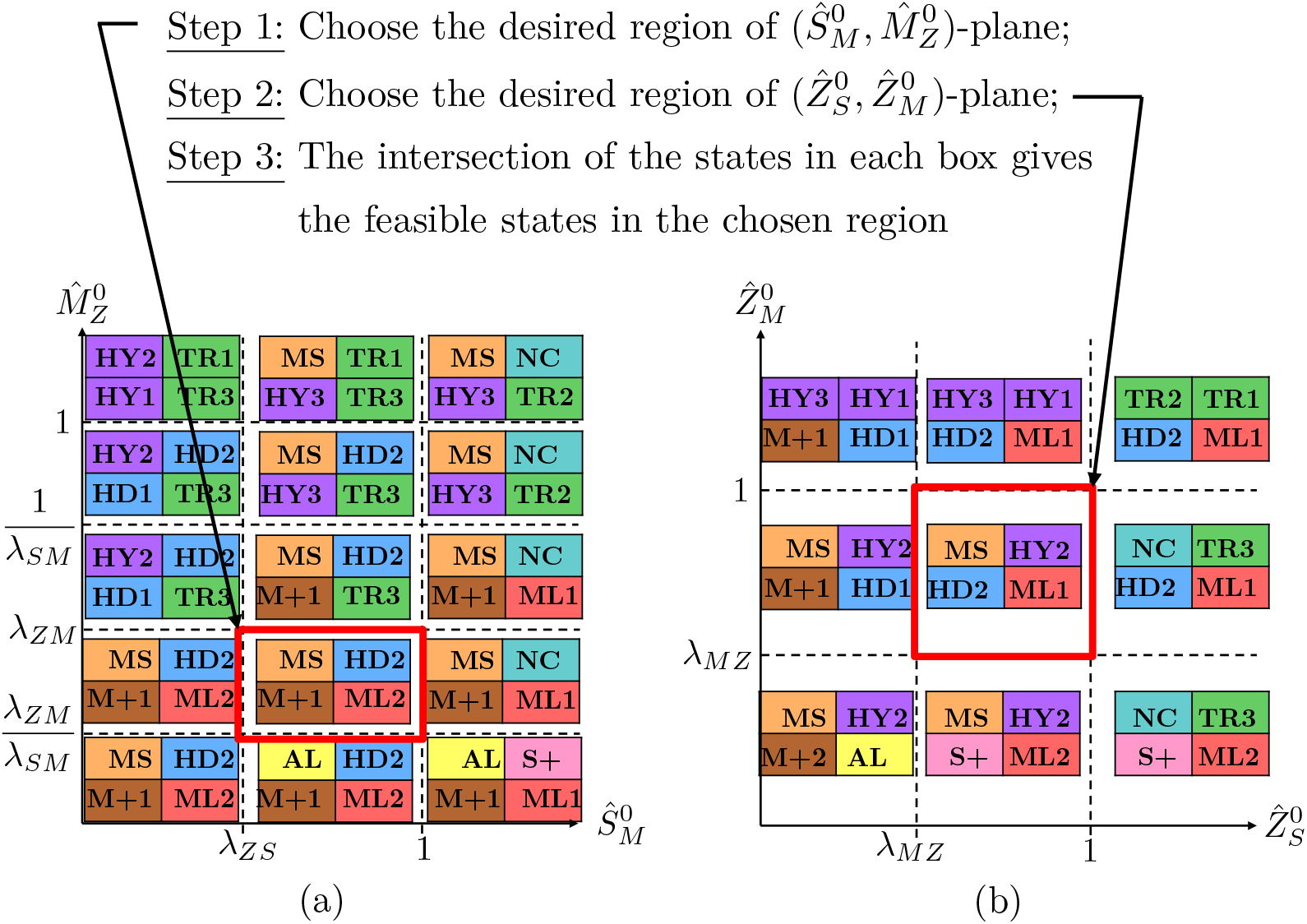
Illustration of spaces spanned by threshold parameters: (a) segments the 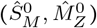-plane; (b) segments the 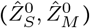-plane. A region of parameter space is constructed by coupling regions from (a) and (b), and feasible temporally stable states are the intersection of states in each box. In the region defined by the red boxes, for example, states MS and HD2 are possible. We exclude saddle points from this Figure.

### 2.3 Simulations

Focusing on bistable parameter regions, we use the tissue-level model to study how subpopulations of cells interact when they are initialised at different locally stable states. We run simulations of the 2D spatial system (7), fixing parameters in a bistable parameter region Ω in which a pair of uniform states **U**_1_ and **U**_2_ (from 1–16 in Table 3) coexist. We initialise a number of clusters in state **U**_1_ by randomly selecting cluster centres and assigning all cell centres within a radius *r* of the seeded centre to state **U**_1_, and the remainder of cells at state **U**_2_. We solve the system as described in Appendix A.5.

We also perform simulations of the one-dimensional (1D) case of a line of cells with uniform spacing and periodic boundary conditions. Fixing parameters in Ω, we perform simulations as described in Section A.5. Taking *N*_*c*_ *=* 100, we initialise cells 33 to 66 at state **U**_1_ and the remainder of cells at state **U**_2_. We explore whether one of the pair of stable states will be preferred by the system when spatial heterogeneity is introduced, testing the spatiotemporal stability of states in Table 3. Simulations are reported in Sections 3.2 and 3.3.

### 2.4 Travelling-wave analysis

We explore the existence of travelling-wave solutions to (7) in the 1D case of a line of cells with uniform spacing and periodic boundaries. In particular, we seek travelling-wave solutions between a pair of stable uniform states from the list in Table 3. Analysis throughout this section is for a bistable parameter region which we call Ω, where some pair of stable states **U**_1_ = (*S*_1_, *M*_1_, *Z*_1_) and **U**_2_ = (*S*_2_, *M*_2_, *Z*_2_) are feasible.

We take the space-continuous limit by assuming 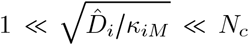, for all *i* ∈ {*S, M, Z*} (taking *κ*_*MM*_ = 1), ensuring that the characteristic wave widths in *Ŝ*, 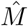 and 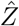 are contained in the domain and are resolved by the grid spacing, i.e. the wavefront spans multiple cells. In this limit, we do not expect the wavespeed to be affected significantly by the periodic boundary conditions. Changing variables to a moving coordinate frame, we uncover a system of ODEs whose uniform states correspond to travelling-wave solutions of (7). Let 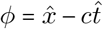, where 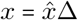 for cell width Δ, *x* is the spatial coordinate and *c* is the speed of the travelling wave. We seek smooth functions *s*(*ϕ*), *m*(*ϕ*) and *z*(*ϕ*) approximating *Ŝ*, 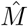 and 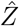 of the form 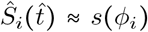 where 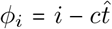. We consider solutions defined on *ϕ* ∈ (−∞, ∞), with boundary conditions (*s, m, z*)(*ϕ*) = **U**_1_ as *ϕ* → −∞ and (*s, m, z*)(*ϕ*) = **U**_2_ as *ϕ* → ∞. Using 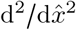 as the space-continuous approximation of the discrete Laplacian 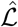 [30], we reformulate (7) in the limit *n* → ∞ as

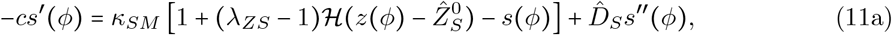

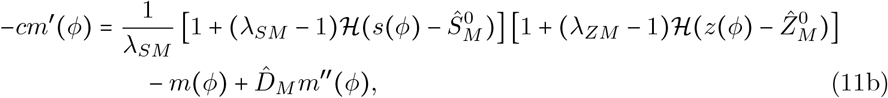

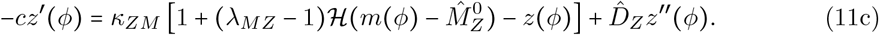

We introduce variables *v*_*s*_ (*ϕ*) ≡ *s*′ (*ϕ*), *v*_*m*_ (*ϕ*) ≡ *m*′ (*ϕ*), *v*_*z*_ (*ϕ*) ≡ *z*′ (*ϕ*) to rewrite (11) as a 6-dimensional system of first-order ODEs (Appendix B.1).

The uniform states of (11) align with the uniform steady states in Table 3. In a bistable region of parameter space, writing **u**(*ϕ*) ≡ (*s, m, z, v*_*s*_, *v*_*m*_, *v*_*z*_)^⊤^, we write the two steady states as **u**_*i*_ = (*S*_*i*_, *M*_*i*_, *Z*_*i*_, 0, 0, 0)^⊤^ for *i* = 1, 2. The Jacobian of the ODE system (B15) can be written in block matrix form as

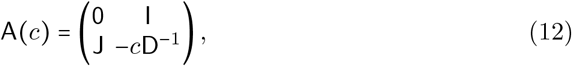

where 0 and I are the 3 × 3 zero and identity matrices, respectively; J is the Jacobian defined in (A11); and 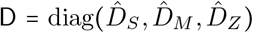. Solving det(A − *µ*I) = 0 at **u**_1_ and **u**_2_ results in the characteristic equation

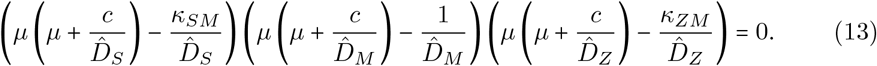

The eigenvalues of states **u**_1_ and **u**_2_ coincide, and are

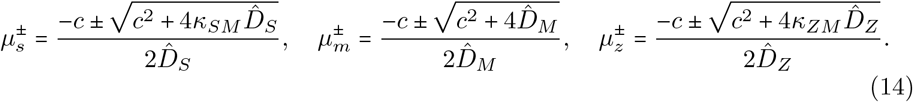

All eigenvalues 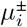 are real with 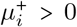 and 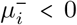 for *c* > 0, and 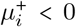 and 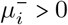 for *c* < 0, for *i* ∈ {*s, m, z*}. Therefore **u**_1_ and **u**_2_ are saddle points in the moving coordinate frame. The corresponding eigenvectors of (14) are

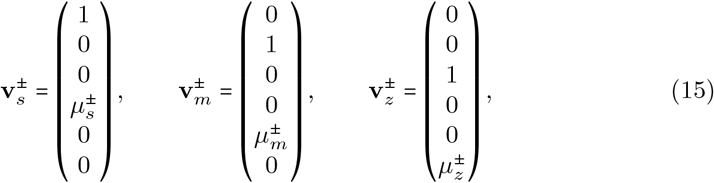

where (for *ϕ*) 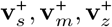 span the unstable manifold and 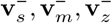 span the stable manifold of each saddle point. A travelling wave from state **u**_1_ to **u**_2_ in the moving coordinate frame takes the form

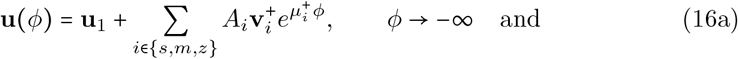

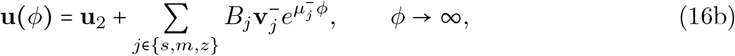

where *A*_*i*_, *B*_*j*_ are constants to be determined.

There are four threshold values (namely 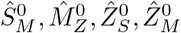) at which each of the four Heaviside functions in (11) switches from one value to another. We construct a piecewise-exponential equation for the travelling wave, which changes when each of these boundaries is crossed. We denote the *ϕ*-values at which these switches occur by 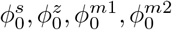, such that

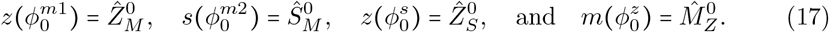

In order to construct the full piecewise-exponential solution, we consider the effects of the changes in each Heaviside function across all variables. We use the boundary 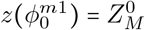 as an example. As this boundary is crossed by the solution, 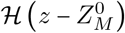 changes value from 0 to 1. Meanwhile, 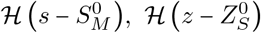, and 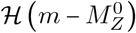 remain constant. Therefore, only the *m* and *m′* (second and fifth) components of the solution will change. We see the same as each boundary is crossed; only one Heaviside function changes value at a time (assuming threshold values are distinct): see Table B4. We use this to reduce the number of unknown coefficients, and write

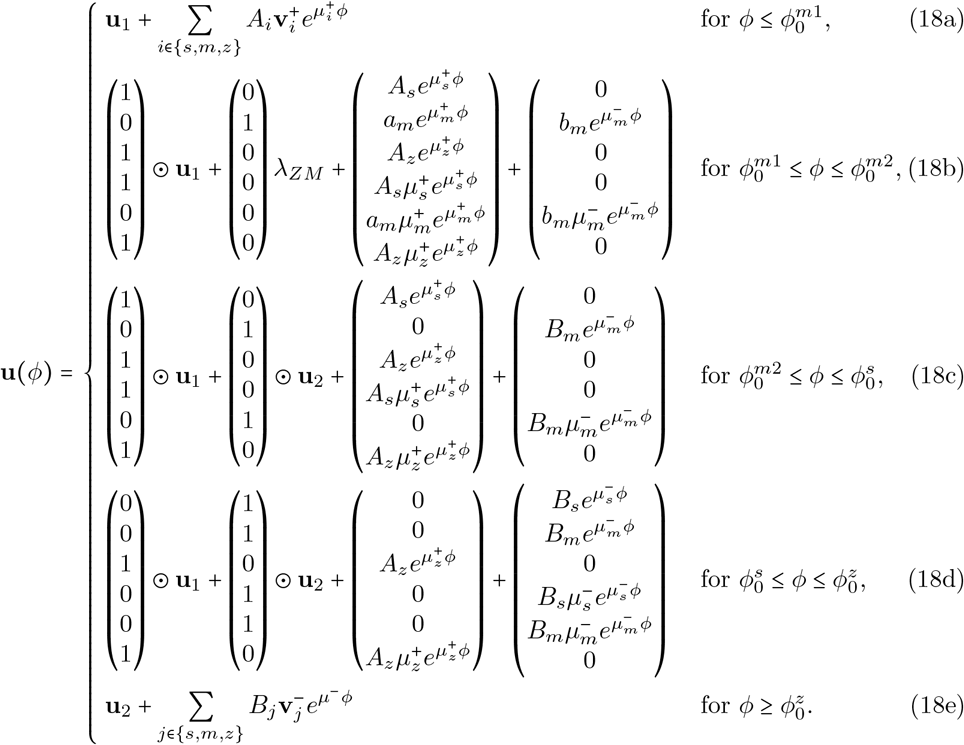

Here, for **u, v** ∈ ℝ^6^, **u** ⊙ **v** = (*u*_1_ *v*_1_, …, *u*_6_ *v*_6_)^⊤^ is the Hadamard product. We impose continuity and smoothness boundary conditions: **u**(*ϕ*) must be continuous at 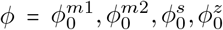 (such that *s, m, z* and their derivatives agree on either side) and (17) must be satisfied. For illustration we have taken 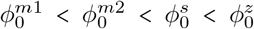; see Appendix B.3 for other cases.

Rearranging the eigenvalue expressions (14) gives

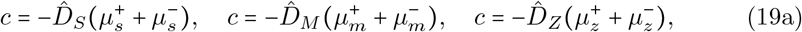

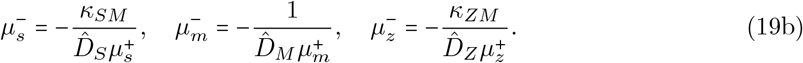

Taking 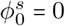 (without loss of generality) and using boundary conditions, we uncover 17 simultaneous equations with 17 unknowns (namely 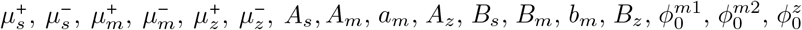), detailed in (B16). We then consider the case *c* =0, a stationary wave. The dispersion relations (19) imply 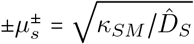, and 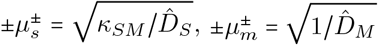. We will call these *µ*_*s*_, *µ*_*m*_, *µ*_*z*_, respectively. In this case the system is over-determined, and solving (B16) results in the condition

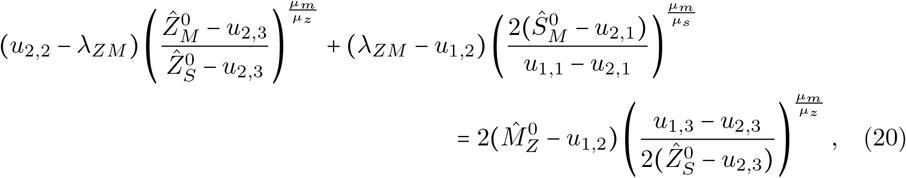

where *u*_*i,j*_ = {**u**_*i*_}_*j*_. The surface (20) separates Ω into two regions, in which *c* < 0 or *c* > 0. If LHS > RHS in (20), **u**_2_ is the spatiotemporally-stable state; if LHS < RHS, **u**_1_ is the spatiotemporally-stable state. If 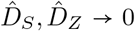, equation (20) reduces to 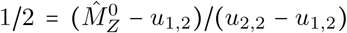, implying that the spatiotemporally-stable state is determined by the distance of each state to the saddle point in the 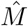-plane (see SA1–SA3 in Table 3). This is similar to results seen in 1-dimensional models with bistability of uniform steady states [31]. However, (20) has been derived under the assumption 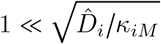; we must therefore assess the predictive accuracy of (20) against simulations.

## 3 Results

In Section 3.1 we classify the steady uniform states in Table 3 in a biologically relevant way and interpret the parameter space illustration from Figure 2 in terms of bifurcations. In Section 3.2 we present spatially-varying simulations in 2D and 1D for an illustrative example of parameter space. We analyse the 1D case by considering travelling-wave solutions in Section 3.3.

### 3.1 Steady uniform states

In the limit *n* → ∞, the full system (7) has up to three steady uniform states, for any given parameter values. Of the possible states, 16 are stable to spatially-uniform perturbations. Under the parameter constraints (10) we label steady-state values as L/M/H in Table 1, indicating that each of the TFs has low/medium/high expression in the cell state, respectively. In Table 2 we formalise phenotypic cell-state definitions into combinations of L/M/H expression across all three TFs. For example, the hyper-differentiated state is defined as any state with MITF being highly expressed and ZEB1 under expressed. The stable states in Table 3 are labelled according to these definitions.

We define the following parameter region, which we will use throughout as an illustration of bistability (the highlighted triangle in Figure 3(c)):

**Fig. 3.**
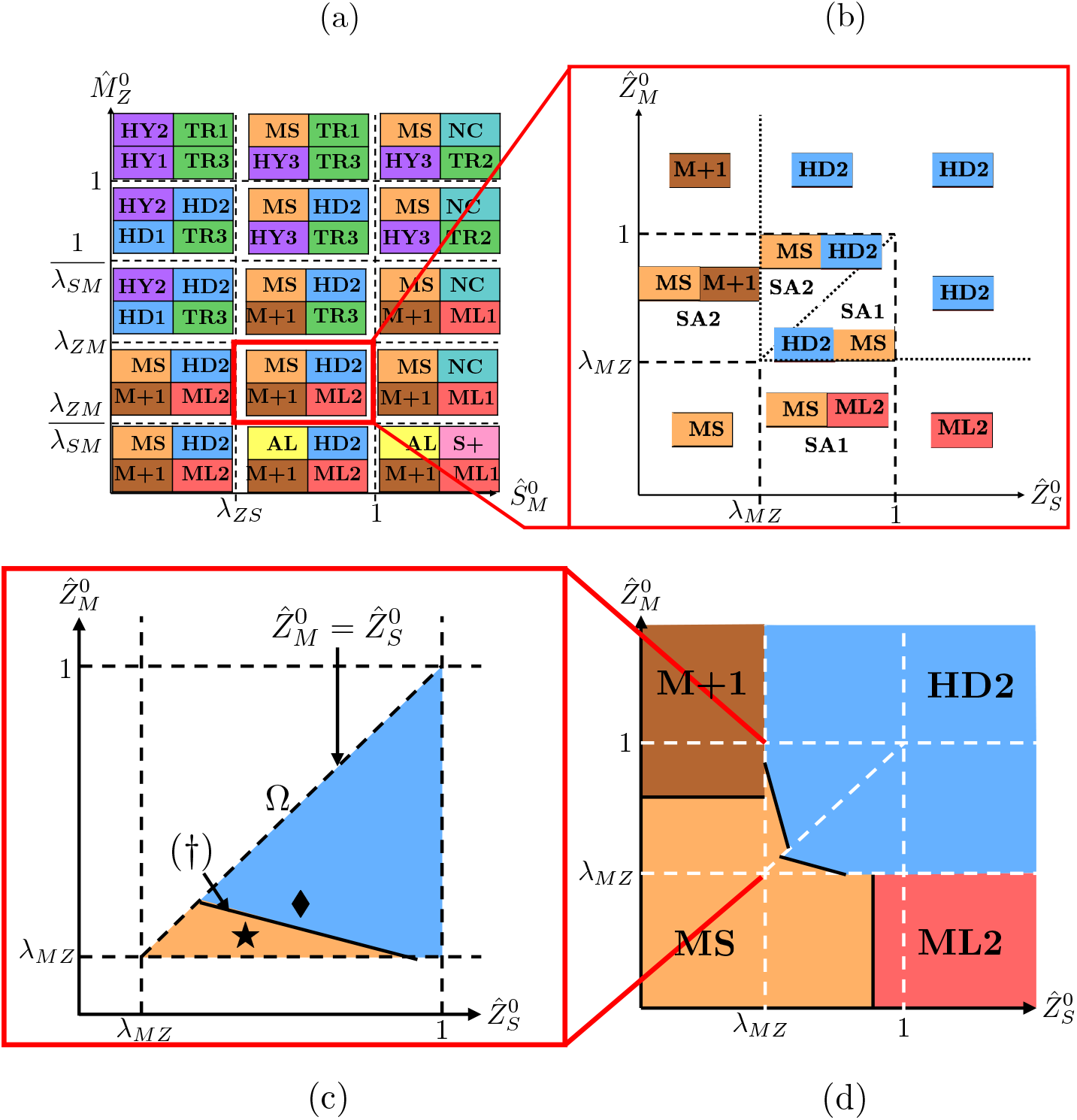
Illustration of solution structure across parameter space for fixed 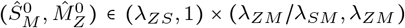. Panels (a) and (b) respectively highlight regions of interest in the 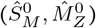 and 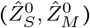 planes. In the highlighted region of 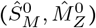-space, saddle-node bifurcations occur at the boundaries of regions labelled SA1 or SA2 in the 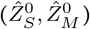 plane. In (a,b), dashed and dotted lines identify thresholds demarcating steady states of different type, as classified in Table 3. In (b), saddle-node bifurcations occur along the dashed boundaries, marking transitions between monostability and bistability. Dotted lines denote changes in the stable state due to changes in value in the Heaviside functions. Panels (c,d) depict regions of spatiotemporal stability of the temporally-stable states in panel (b). Panel (c) depicts the parameter region Ω (21), and the regions of spatiotemporal stability of states MS and HD2; (†) labels the spatiotemporal stability threshold (22); (★)and (⧫) label the parameters used in simulations shown in Figures 4, 5 and 6, 8 respectively, found below.

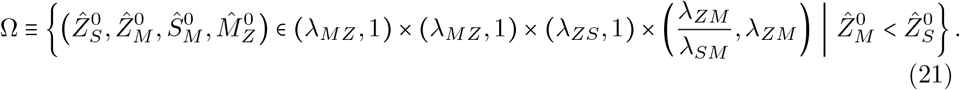

Stable states HD2 and MS coexist in Ω, as well as the unstable saddle point SA1. We fix 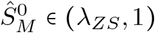 and 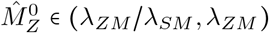 (Figure 3(a)) and illustrate in Figure 3(b) the corresponding parameter regions in the 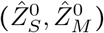-plane. As parameters 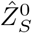 or 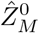 vary, bifurcations occur in which a saddle-node pair is created or annihilated. As parameters move from a bistable region (containing two coloured states and a saddle in Figure 3(b)) to a monostable region (containing a single coloured state), the system is driven towards a single remaining stable state. These saddle-node bifurcations occur along the dashed lines in Figure 3(b). We see a different type of state change along dotted lines. Moving across these boundaries, the system remains monostable. The change in state does not correspond to a bifurcation, but to a discontinuous shift in the same equilibrium branch, caused by changes in the intersection structure of the Heaviside functions (9). This happens as threshold values 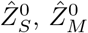 vary, according to the conditions listed in Table A1. In this sense, states M+1 and ML2 are ‘extensions’ of the HD2 solution, part of the same equilibrium branch.

We further explore the relationship between stable uniform states, and their spatiotemporal stability, through simulations of the tissue-level model in the bistable parameter region depicted in Figure 3(c).

### 3.2 Tissue-level simulations in 2D

We fix parameters in Ω (21), highlighted with the marker (⧫) in Figure 3(c). Figure 4 shows snapshots of simulations of (7) over an array of cells (as described in Section 2.3) for varying diffusion coefficients 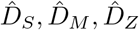. We initialise clusters of cells at state HD2 and the remainder of cells at state MS; in Figure 4(a-c), where 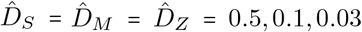, respectively, state HD2 dominates the population. This indicates that HD2 is spatiotemporally-stable and MS is not. In Figure 4(d), where 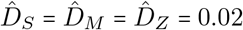, we see an initial invasion of state HD2 into state MS, however the invasion eventually stops, leaving a cluster of MS cells that persist within the HD2 population.

**Fig. 4.**
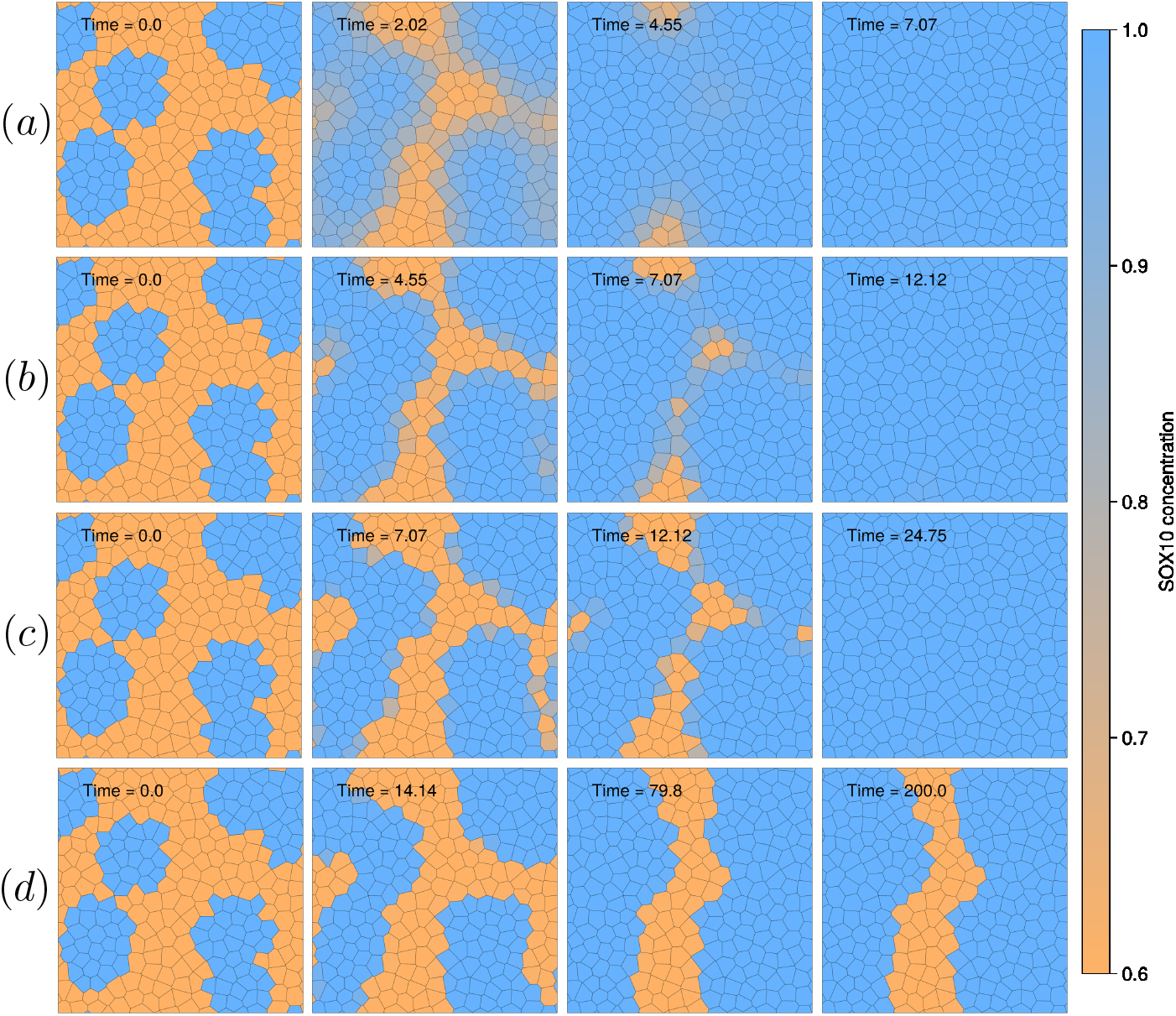
Snapshots from simulations using (7) of the 2D disordered system (see Appendix A.2), for varying diffusion coefficients, 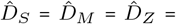 (a) 0.5; (b) 0.1; (c) 0.03; (d) 0.02. Values of 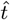 are annotated for each snapshot. We use SOX10 expression as a proxy for the cell state. Blue cells are in state HD2 and orange cells in state MS (Table 3). Parameters used are: *N*_*c*_ 253, *λ*_*SM*_ 5, *λ*_*MZ*_ = 0.3, *λ*_*ZS*_ = 0.6, *λ*_*ZM*_ = 0.1, 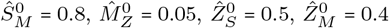 (in Ω (21), marked with a diamond in Figure 3(c)), *K*_*S*_ = *K*_*M*_ = *K*_*Z*_ = 1 and *L*_*x*_ = *L*_*y*_ = 10.

### 3.3 Tissue-level simulations in 1D

We perform simulations for the same set of parameters in the 1D case, taking 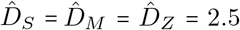, to gain further insight into this behaviour (Figure 5). We initialise 16 cells 33 to 66 (in a ring of 100 cells) in state HD2 and the remainder of cells in state MS. State HD2 ultimately dominates the population of cells (Figure 5). When we alter parameters 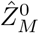 and 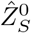 (while remaining in Ω, highlighted by the marker (★) in Figure 3(c)), state MS ultimately dominates the line of cells (Figure 6). In both cases, we verify the existence of travelling-wave solutions by tracking points in *Ŝ*, 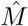 and 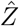 over time. We estimate wave speeds directly from the simulations in Figures 5 and 6 (as detailed in Appendix A.5), with *c* ≈ 0.9 in Figure 5 and *c* ≈ −0.2 in Figure 6, where the sign is associated with the wave in the right-hand part of the domain. For all *c >* 0, state HD2 will dominate the system and for *c* < 0 state MS will dominate the system.

**Fig. 5.**
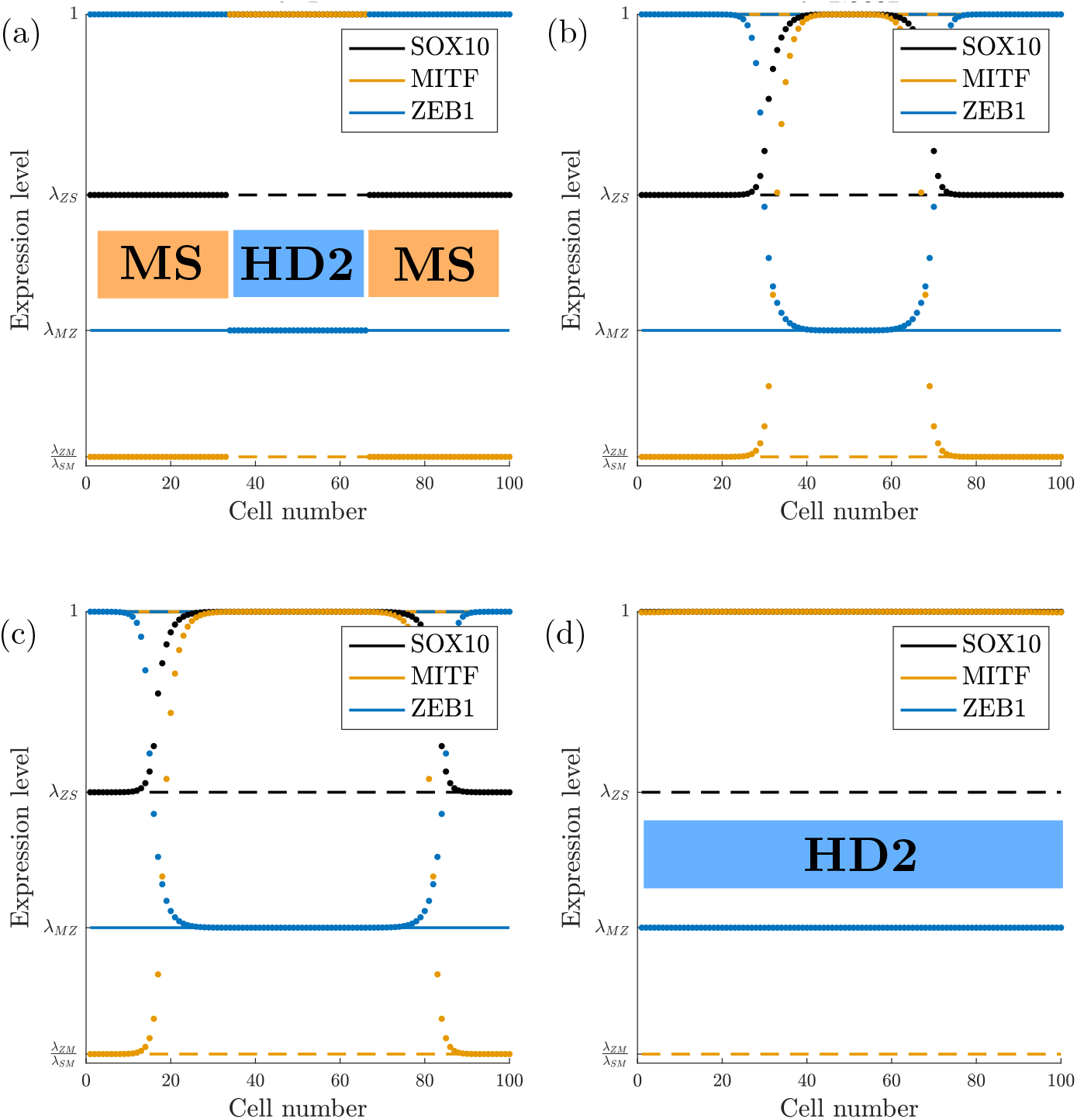
Snapshots from a simulation of the 1D spatiotemporal system (7) (*N* = 100) with periodic boundaries, as described in Appendix A.5, at (a) 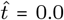 (b) 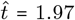 (c) 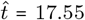 (d) 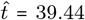 Circular plots are values of *S,M, Z* in each cell (along the horizontal axis). Solid/dotted lines represent *S,M, Z* levels at steady states HD2/MS, respectively. We take 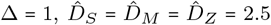, *λ*_*SM*_ = 5, *λ*_*MZ*_ = 0.3, *λ*_*ZS*_ = 0.6, *λ*_*ZM*_ = 0.1, 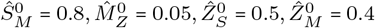 (in Ω (21), (marked with a diamond in Figure 3(c) and *K*_*S*_ = *K*_*M*_ = *K*_*Z*_ = 1.

**Fig. 6.**
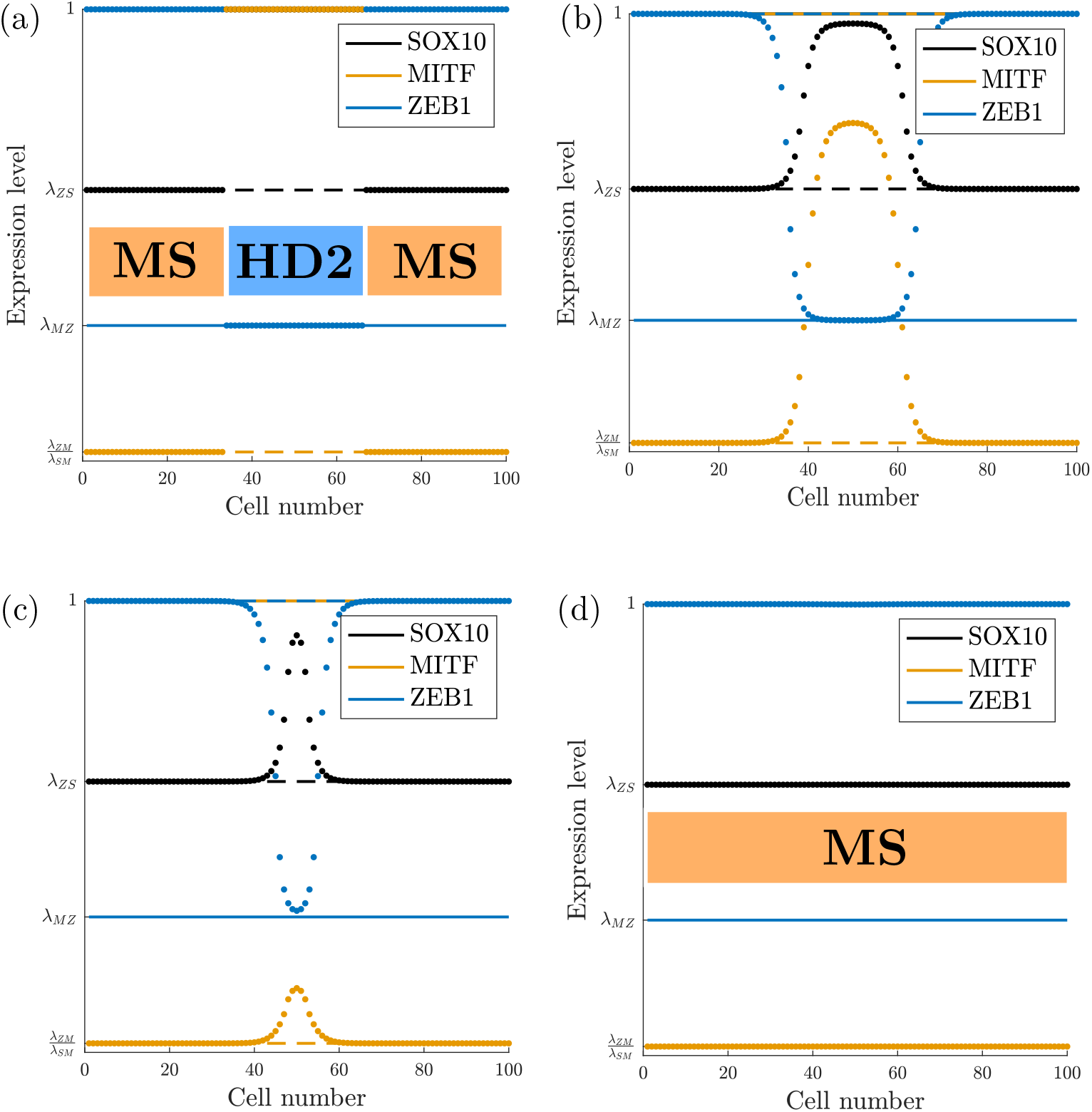
Snapshots from a simulation of the 1D spatiotemporal system (7) (*N*_c_ = 100) with periodic boundaries, as described in Appendix A.5, at (a) 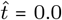 (b) 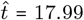 (c) 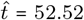 (d) 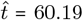. Circular plots are values of *S,M, Z* in each cell (along the horizontal axis). Solid/dotted lines represent *S, M, Z* levels at steady states HD2/MS, respectively. We take 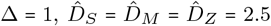, *λ*_*SM*_ =5, *λ*_*MZ*_ = 0.3, *λ*_*ZS*_ = 0.6, *λ*_*ZM*_ = 0.1, 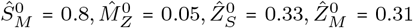 (in Ω (21), marked with a star in Figure 3(c) and *K*_*S*_ = *K*_*M*_ = *K*_*Z*_ = 1.

Figure 7 overlays simulations in Figure 5 with the travelling-wave solution (18) for the chosen parameters, where exponential coefficients are found by solving the simultaneous equations (B16) numerically. Numerical predictions of *c* from (B16) (*c =* 0.88 for the simulation in Figure 5, *c* = −0.23 for Figure 6) match well with estimates from tracking a point over time on each wave in simulations (Appendix A.5). We label with coloured stars the points at which the piecewise exponential solution (18) switches, which corresponds to when each variable crosses its Hill threshold value. The space-continuous limit gives a good approximation of the travelling wave, so we use this to study the direction of the wave in the discrete system.

**Fig. 7.**
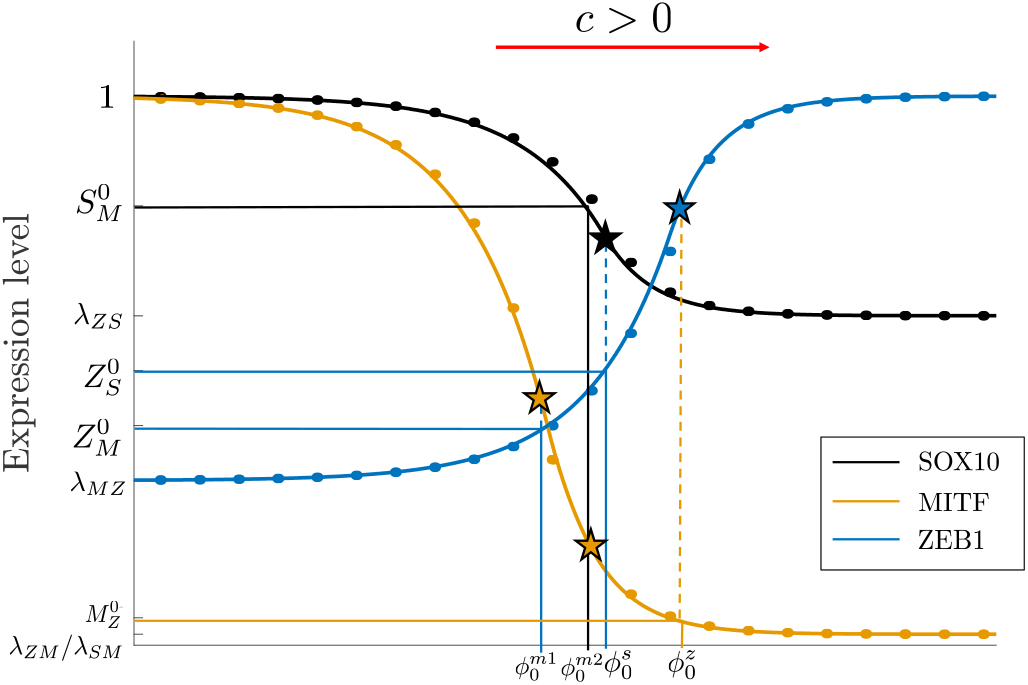
Figure showing a comparison between an overlay plot of simulations (circular markers) and the analytical solution (18) (solid lines, using the coefficients as solutions of (B16)), and an illustration of the switching points of each exponential. Simulation plots are overlay plots of the right-moving wave seen in Figure 5, at 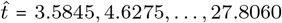. Snapshots are translated with *c* 0.9 (see A.5). Coloured stars highlight the points along the wave where the exponentials in the continuous travelling-wave solution (18) switch.

In the simulations in Section 3.2, we take *K*_*S*_ = *K*_*M*_ = *K*_*Z*_ and 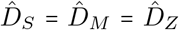,giving 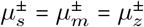. Under these parameters, the surface (20) has the form

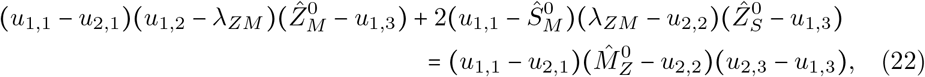

where HD2 = (*u*_1,1_, *u*_1,2_, *u*_1,3_)^⊤^, MS = (*u*_2,1_, *u*_2,2_, *u*_2,3_)^⊤^ and 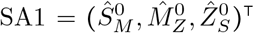. This surface divides Ω into two regions in which MS or HD2 will be the dominant state across the line of cells. The projection of (22) onto the 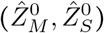-plane (for fixed 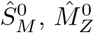) is illustrated in Figure 3(c) as a straight line through Ω, labelled (†).

We label the left- and right-hand sides of (22) as LHS and RHS, respectively. If LHS > RHS (i.e., if parameters lie in the blue region of Figure 3(c)) then *c* > 0 and HD2 is the spatiotemporally-stable state, while if LHS < RHS (i.e., if parameters lie in the orange region of Figure 3(c)) then *c <* 0 and MS is the spatiotemporally-stable state, consistent with simulations in Figures 5 (⧫) and 6 (★). In Figure 8, we run simulations over the 2D monolayer for the same parameters as the simulations in Figure 6 varying 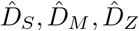, for which LHS < RHS in (22). In Figure 8(a) and (b), where 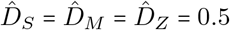 and 0.1, respectively, state MS invades the population, in agreement with the 1D stability prediction for these parameters. In Figure 8(c) and (d), where 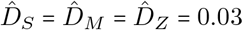 and 0.02, respectively, we observe similar behaviour to Figure 4(d), with state MS initially invading before the invasion stops (particularly in Figure 8(d) where very little change from the initial condition is observed). This suggests that the 1D stability prediction only holds in 2D when the characteristic wavelength (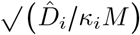 for *i* ∈ {*S, M, Z*}) is sufficiently large relative to the cell size.

**Fig. 8.**
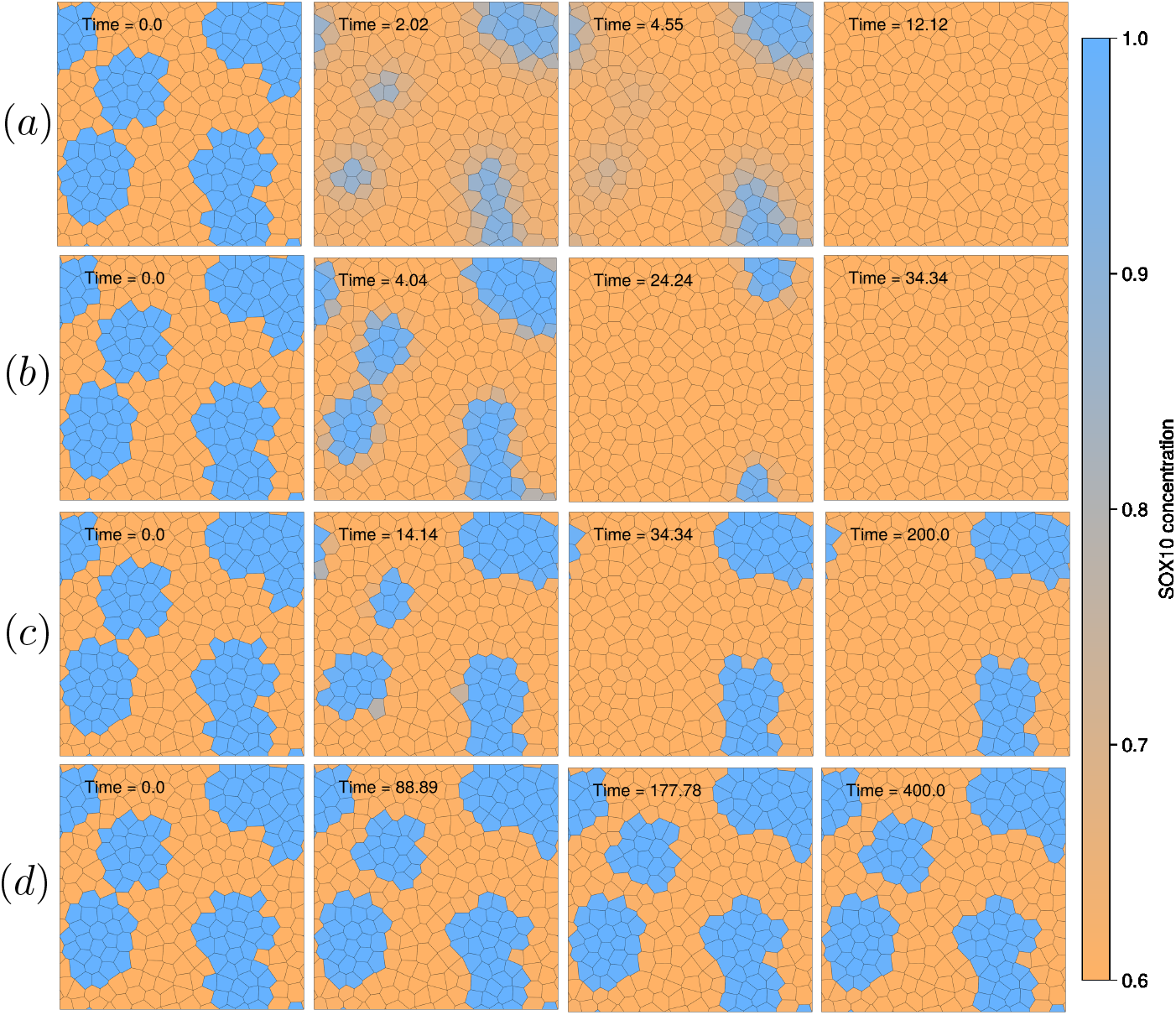
Snapshots of 2D simulations over the same monolayer and initial conditions as Figure 4, with 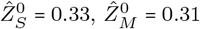 such that LHS < RHS in (22) (marked with a star in Figure 3(c)). Values of 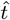 are annotated for each snapshot. Diffusion coefficients are 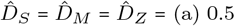; (b) 0.1; (c) 0.03; (d) 0.01.

The condition (22) is consistent with the saddle-node bifurcation that occurs between states MS and SA1 at the boundary of Ω (21) as the bifurcation parameter 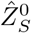 increases through unity. The condition reduces to 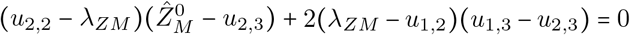. Since all the terms in this expression are positive, LHS > RHS trivially. HD2 is therefore the spatiotemporally-stable state, consistent with Figure 3(c).

Extending the same analysis to the remaining bistable regions in Figure 3(b) yields similar results, illustrated in Figure 3(d). The methodology is given in Appendix B.3, and involves solving the relevant simultaneous equations (cf. (B16)) for the case *c =* 0 to derive the associated spatiotemporal stability thresholds. The thresholds in Figure 3(d) divide 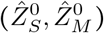-space into two regions: a region in which MS is the spatiotemporally stable state; and a region in which HD2 (and its extensions M+1, ML2) is the the spatiotemporally stable state. The discontinuities in the boundaries in Figure 3(d) arise due to changes in the threshold equation and discontinuous changes in the values of **u**_1_,**u**_2_ between parameter regions.

## 4 Discussion

We have established that the tissue-level model, defined in Section 2, with three interacting species and four thresholds, accommodates 19 steady uniform states under a biologically relevant classification, which we are able to write analytically in the limit *n* → ∞ (assuming large numbers of transcription factors bind to promoter sites). By mapping the steady uniform states onto a four-dimensional projection of the high-dimensional parameter space (Figure 2), we visualise how these states populate different regions of parameter space, defined by varying the threshold values of the GRN. This visualisation reveals that all parameter regions are either temporally monostable or bistable, as well as saddle-node bifurcations between these regions. This bistability is disrupted for a population of cells, with the system choosing one state over another through travelling waves of gene expression, under sufficiently strong cell-cell signalling. We can predict the stable state of a population of cells using system parameters: in a temporally bistable region, a spatiotemporal stability threshold determines which of the two states will be preferred by the system, as illustrated in Figure 3.

We employ a different classification of cell states to Subhadarshini *et al*. [19]. In line with their findings, the GRN allows for a range of cell states, which can determine different phenotypes [4, 6, 7, 16, 19, 32, 33]. Subhadarshini *et al*. classify emergent states from simulations by normalising the values of each variable across all states and hierarchically clustering these values into four groups. They define a proliferative score (the sum of SOX10 and MITF expression) and an invasive score (which is a sum of SOX9, JUN and ZEB1) and plot the simulated steady states on a proliferative-invasive axis. We distinguish between MITF and SOX10 to consider the three TFs separately. We take an empirical stance on the classification of cell states, motivated by experimental observations and the literature [2, 3, 9, 14, 16], and use restrictions on system parameters to label each stable state value as low, medium or high expression in Table 1. This approach enables greater resolution and reveals more subtle differences between cell states than Subhadarshini *et al*., which may be more reflective of the recent literature showing experimental sub-clustering of melanoma cells into upwards of 11 distinct subgroups [16, 33]. We do not impose assumptions about the biological similarity between the cell states in Table 2 and reported biological phenotypes, although some are biologically proximal (e.g. more invasive or proliferative phenotypes). The states presented here have emerged from the mathematical model, and while we assign biologically motivated labels, some may correspond to parameter regions that are not feasible.

Under the assumption that large numbers of TFs bind to promoter sites (*n*→ ∞) we reduce the model of rate of change of TF expression levels to piecewise constant form, making the overall model piecewise-linear and non-smooth. In contrast to [19], this simplification allows us to determine exactly how many steady states the GRN admits, providing a comprehensive understanding of the system. Stability analysis in this limit reveals there are 16 stable states (which we classify as previously described) and 3 unstable saddle points, whose existence would not have been apparent from simulations alone. The large-*n* assumption does not strictly hold in biological systems that require low molecule numbers for cooperative binding. The sharp transitions that we observe will be smoothed out for finite *n*. In Appendix C we explore the temporal bistability behaviour of the system for a range of values of *n* and observe that for *n* ≥ 7 the structures qualitatively resemble those seen in our approximation. For smaller values of *n* we see more complex behaviour, including a butterfly catastrophe arising near *n =* 5 [34] in which pockets of parameter space display tristable behaviour (with three stable states). While this structure is sensitive to small variations in *n*, the features revealed in the large-*n* limit are more robust, making the assumption a useful approximation of the system’s behaviour.

By introducing spatial variations in expression, we study spatiotemporal stability of states across a population of cells via travelling waves in gene expression. In (19) we give expressions for the wave speed in terms of nondimensional diffusion coefficients and eigenvalues. In dimensional form, these can be written

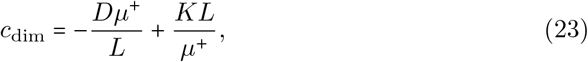

for *D =* {*D*_*S*_, *D*_*M*_, *D*_*Z*_}, 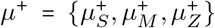 and *K =* {*K*_*S*_, *K*_*M*_, *K*_*Z*_}, respectively, where *L* is the cell length scale defined in Appendix A.2. This shows that the speed of propagation of the travelling wave is mediated both by the diffusion coefficients *D* and by the degradation rates *K* of each TF. The predictions made by the 1D travelling-wave analysis hold only for values of *D* and *K* such that the characteristic wavelength is sufficiently large relative to the cell width. Figures 4(d) and 8(c,d), showing simulations for small diffusion coefficients, reveal an initial propagation before the wave of expression comes to a halt, resulting in a stable pattern of heterogeneous expression across the population of cells. Thus, when neighbouring cells can communicate sufficiently strongly, the bistability in the model is disrupted. Rather, the system selects a single spatiotemporally stable state from the two temporally stable ones. In the example presented in Section 3.3, the system chooses between a hyper-differentiated or mesenchymal-like phenotypic state. In this case, the dominant state can be changed from hyper-differentiated to mesenchymal-like by decreasing the threshold for ZEB1 expression for inhibition of SOX10 and/or MITF, to cross the boundary defined in (22). While we do not have biological evidence of this, the images in Figure 1 do show subpopulations of cells taking on different phenotypic cell states, with the oval regions showing high SOX10, high MITF and low ZEB1 (characteristic of a hyper-differentiated cell state) and other tumour cells exhibiting low SOX10, low MITF and high ZEB1 (characteristic of a mesenchymal cell state). A potential prediction from the model is that these subpopulations lie in different regions of parameter space, or there is a physical block for the travelling wave of gene expression, e.g. via suppression of cell-cell signalling (reminiscent of the patterns in expression in Figures 4(d) and 8(c,d)). By calculating the spatiotemporal stability threshold for various parameter regions, we see in Figure 3 that parameter space is divided into wider regions in which each state is spatiotemporally stable.

The parameter landscape we present leads to some experimental hypotheses. As shown in Figure 3, state transitions occur as the thresholds for inhibition or activation vary, namely 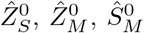, and 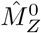. These are nondimensionalised threshold values that reflect the ratio between the production and degradation rates of each TF. So, when a threshold value increases in the nondimensional model, this may correspond to an increase in the underlying production-to-degradation ratio. Endoplasmic Reticulum Associated Degradation is known to be impacted in melanoma [35–37]. Alternately, a change in the threshold of TF activation/repression of a gene could theoretically occur through other mechanisms, for example, a change in the number of available TF binding sites, such as by epigenetic alterations, which open or close chromatin to reveal or obscure binding sites [38]. Some enzymes can biochemically alter chromatin by adding or removing acetylation or methylation groups, which in turn changes chromatin accessibility and accessibility to TF binding sites. These enzyme targets are currently being exploited in the clinic against many cancers [39]. Other causes of a change in threshold levels include mechanical stress (such as compression), which has been shown to induce chromatin changes [40]; mutations in the genome across cells; and extracellular signals [41]. Furthermore, the bifurcation diagram in Figure 3 suggests specific pathways of state transitions. A possible hypothesis from this model is a prediction of which parameter changes drive nevus cells to develop melanoma-like characteristics or identify the parameter regions in which these state transitions occur. Nevus cells may correspond to the hyper-differentiated or MITF^+^ phenotypes in the model, so one might explore parameter regions where these states coexist with a transitory state, a potential pathway for the nevus–to–melanoma transition. The cases we present in Section 3 are illustrative and we do not claim significance for the hyper-differentiated–to–mesenchymal-like (or vice versa) state transition. Identifying common state transitions experimentally could indicate which parameter region a system lies in and could provide validation of these theoretical pathways. For example, if transitions between mesenchymal-like and neural-crest like states were more frequently observed, the model would be tested in parameter regions where such states are predicted to exist.

The travelling-wave analysis we present suggests that intercellular communication via diffusing signalling molecules may contribute to gene expression patterns in a tumour, such as those observed in Figure 1, where subpopulations of cells share a common phenotypic state. If it is possible to identify a diffusing signal experimentally, the model suggests perturbing this signal could alter the observed pattern. In the simulations shown in Figures 4 and 8, we see that for larger diffusion coefficients a single cell state is adopted by the population, whereas for smaller diffusion coefficients there is heterogeneity in the stable states across the population. Phenotypically homogeneous tumour regions may therefore exhibit stronger cell-cell communication, or involve a diffusing signal with a longer effective range. Gopalan *et al*. [16] observe an increase in regions of phenotypically homogeneous tumour cells (so-called ‘encapsulated’ regions) after anti-PD-1 therapy in mouse melanoma models, within which cells are more differentiated (i.e., cells express high levels of the MITF-induced gene, Rab38). This may imply that communication is stronger after therapy, particularly within these encapsulated regions. Eom *et al*. [42] see similar spatial patterns in human lymph nodes infiltrated with melanoma, with homogeneous clusters of MART1 expressing differentiated cells.

The methods presented in this paper depend on parameters whose values are not precisely known. Although this introduces uncertainty, these results frame discussions about the unknown parameters. Subhadarshini *et al*. [19] provide biologically relevant ranges for GRN parameters. Various other experimental and modelling studies report GRN parameter values, for different proteins and regulatory networks, but give an impression of the ranges of values of production rates (of order 10^−4^nM/s), degradation rates (of order 10^−4^/s), and diffusion lengths and timescales of signalling molecules whose values can vary between 100−150*µ*m and 5−90 minutes, respectively, for sensing cells with a similar cell width to those in our system [43–46]. From the scale bar in Figure 1, we estimate cell widths to be 10 − 15*µ*m. A direct continuation of this work lies in deriving these parameter values experimentally for this model, which will refine parameter space to the biologically feasible regions.

To focus on core expression dynamics, we have stripped the GRN to only three TFs. In reality, this network will consist of many more genes and TFs, but the interactions used in the GRN are well established, and cell states can be effectively classified using these three. If, for example, we incorporate self-activation for each TF in the GRN, as seen in [19], the number of stable steady states increases by a factor of eight, and parameter space becomes more complex due the introduction of associated threshold and fold change parameters, complicating parameter exploration. This further motivates our focus on a simplified GRN that captures the core network dynamics while remaining mathematically tractable. Other regulatory effects are treated as being absorbed into the production and degradation rates, or fold changes in expression due to activation and inhibition. Furthermore, we model intercellular communication via diffusion, which is a common approach but a simplification of the complex signalling processes involved in reality, such as receptor-ligand interactions or signal transduction pathways. We do not suggest that TFs physically move between cells; rather, we use diffusion as a mathematically tractable approximation of intercellular communication, providing insight into collective cellular behaviour.

In summary, by assuming that large numbers of transcription factors bind to promoter sites, we have provided a comprehensive analysis of a nonlinear regulatory network that is central to the development and progression of melanoma, classifying all possible phenotypic states and highlighting distinctions in stable gene expression between individual cells and cell populations. We have shown that one phenotype can dominate a population, provided intercellular communication is sufficiently strong. The model provides a foundation for future studies that can incorporate more refined descriptions of the genetic pathways driving malignancy and the spatial organisation of cell populations in a tumour.

## Declarations

### Funding

This work was supported by the British Skin Foundation Young Investigator Award, 023/YI/22, The Wellcome Trust (204796/Z/16/Z) and The University of Manchester FBMH Dean’s Prize award, all awarded to KLM; Cancer Research UK (RCCCDF-Nov24/100004), awarded to KLM, supported KLM, RL and components of this study; Wellcome Trust Career Development Award (225408/Z/22/Z) to SW; CTB was supported by an EPSRC Doctoral Training Studentship.

### Conflict of interest

None

### Ethics approval and consent to participate

Not applicable

### Consent for publication

For the purpose of open access, the authors have applied a Creative Commons Attribution (CC-BY-4.0) licence to any Author Accepted Manuscript version arising.

### Data availability

Not applicable

### Materials availability

Not applicable

### Code availability

Simulations of the 1D model, wave speed and parameter estimation (Section 3.3, B.2) at github.com/charlietaylorbarca1/GRN_Simulations.git. Construction of the disordered monolayer of cells (Section 3.2) at github.com/charlietaylorbarca1/Monolayer_Construction.git. 2D simulations (Section 3.2) at github.com/charlietaylorbarca1/Reaction_Diffusion_Simulations.git. Continuation bifurcation boundaries (Section C) at github.com/charlietaylorbarca1/Bifurcation_Plots.git.

### Author contribution

Conceptualization: CTB, VG, SW, KLM, GWJ, OEJ; Methodology: all authors; Formal analysis and investigation: CTB; Writing - original draft preparation: CTB; Writing - review and editing: all authors; Funding acquisition: SW, KLM.

## Appendix A Model set-up

### A.1 Reaction-based derivation of local GRN model

We build a set of reactions for the GRN (2), first for reactions affecting *Z* as a result of the inhibiting relationship with *M* :

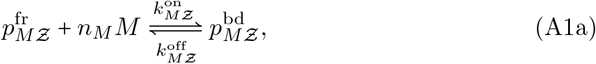

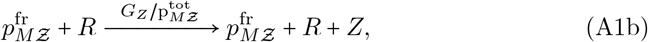

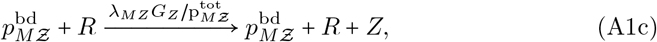

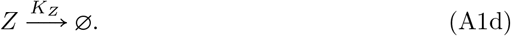

Reaction (A1a) describes the reversible binding of *n*_*M*_ *M*-molecules to free sites on *Ƶ*’s *M*-promoter region with rates 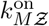 and 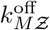. Reaction (A1b) describes the production of TF *Z* due to a free *M*-promoter space; reaction (A1c) describes the production of TF *Z* due to a bound *M*-promoter space. Reactions (A1b) and (A1c) involve an RNA polymerase molecule, *R*, which catalyses the transcription process. *G*_*Z*_ is the maximal production rate of TF *Z* by gene *Ƶ*; the maximal production rate due to an individual promoter is 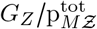, where 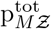 is the total number of spaces on *Ƶ*’s *M*-promoter. The factor *λ*_*MZ*_ *<* 1 in (A1c) models the way in which occupied promoter spaces restrict the production of *Z*. Finally, (A1d) describes the degradation of *Z*, with a degradation rate constant *K*_*Z*_.

We build a similar set of reactions that affect *S*, as a result of the inhibiting relationship with *Z*:

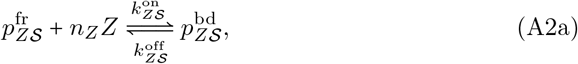

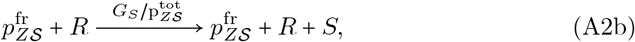

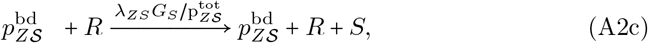

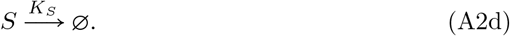

We consider two separate promoter regions on ℳ, the *Z*- and the *S*-promoter, and how combinations of bound/free *Z*- and *S*-promoter spaces will collectively affect the production rate of *M* :

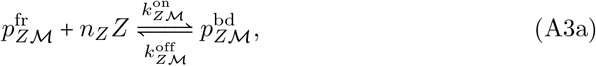

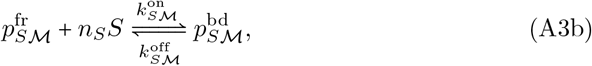

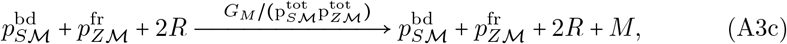

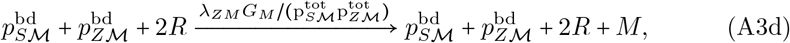

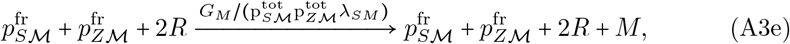

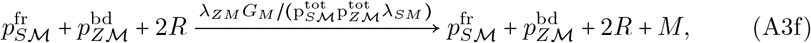

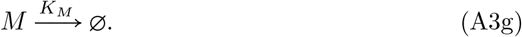

*G*_*M*_ is the maximal production rate of *M*, which is achieved by the combination of a bound *S*-promoter space and a free *Z*-promoter space. Bound *Z*-promoters will restrict production as in the previous cases, with *λ*_*ZM*_ *<* 1, and free *S*-promoters will achieve slower production, with *λ*_*SM*_ > 1. The 2*R* terms in reactions (A3c)–(A3f) describe separate RNA polymerases binding to the distinct promoter regions.

We describe reactions (A1)–(A3) as a system of coupled ODEs using mass action kinetics. We use the following notation: *S* = [*S*], *M* = [*M*], *Z* = [*Z*], and 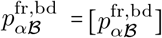 for *α* ∈ {*S, M, Z*}, ℬ ∈ {𝒮, ℳ, *Ƶ*} and use

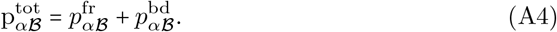

to specify 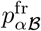 in terms of 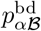. We take the concentration of RNA polymerase in the nucleus to be constant, *R*, and absorb these constants into the reaction rates. The ODE system that describes the GRN according to reactions (A1)–(A3) is

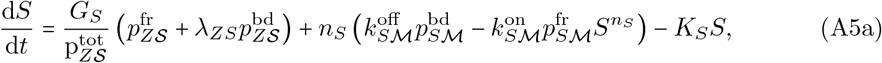

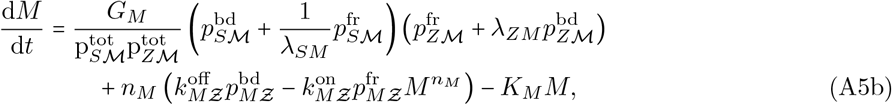

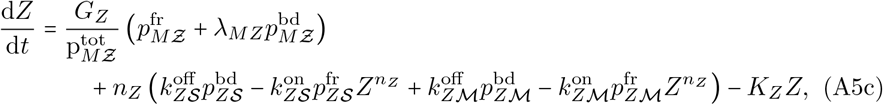

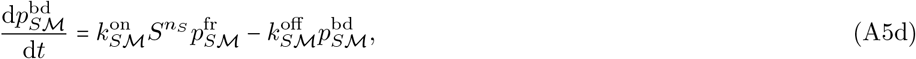

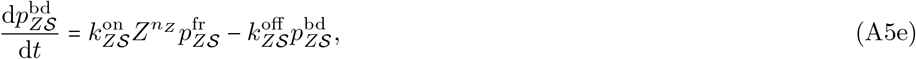

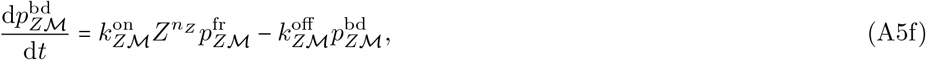

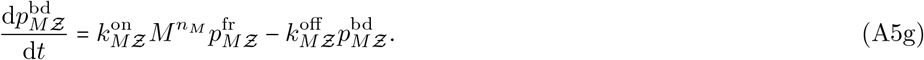

We assume that the binding and unbinding processes of TFs from promoter regions are in approximate equilibrium. We therefore simplify the system of ODEs (A5a) using the quasi-steady-state hypothesis 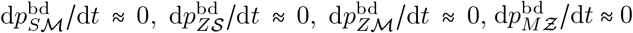, under which, and using (A4), we obtain

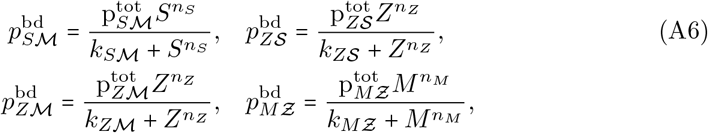

where 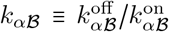. Taking *n*_*S*_ = *n*_*M*_ = *n*_*Z*_ = *n*, say, we set 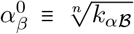, which can be interpreted as the ratio of binding-unbinding rate constants for each molecule involved in cooperative binding. Substituting (A6) into (A5a), and once again using (A4), results in the system (4).

### A.2 Spatiotemporal model

We adopt the notation and tools used in [30] for diffusion over a disordered monolayer of cells. Cell centres are seeded in a periodic box of size *L*_*x*_ × *L*_*y*_ through a Matérn process and connected through Delaunay triangulation. We use periodic boundaries such that centres can be connected across boundaries. We assign a region to each cell using Voronoi tessellation, depicted in Figure A1. We will refer to the Voronoi network as the primal network, and to the Delaunay network as the dual network. We label vertices, faces and edges of the primal network by *k* ∈ {1, …, *N*_*v*_}, *i* ∈ {1, …, *N*_*c*_} and *j* ∈ {1, …, *N*_*e*_}, respectively.

Cell edges and faces are assigned an orientation and we define signed incidence matrices A and B as follows [47]. For an edge *j* exiting *k*′ and entering *k*′′,

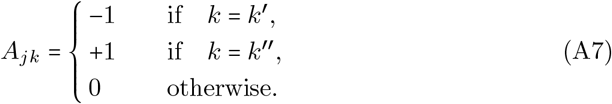

*B*_*ij*_ = 1 if edge *j* is a part of cell *i* and has congruent orientation, *B*_*ij*_ = −1 if edge *j* is a part of cell *i* and has opposing orientation, and *B*_*ij*_ = 0 otherwise. Ā and 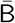 are the unsigned incidence matrices. Vertices on the primal network are at **r**_*k*_ ∈ *ξ*, where *ξ* is the domain depicted in Figure A1. We set edge vectors and lengths **t**_*j*_ = ∑_*k*_ *A*_*jk*_**r**_*k*_, *t*_*j=*_ |**t**_*j*_∣, and edge centroids 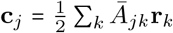 (with adjustments made for cells on the boundary). We define cell centres (vertices of the dual network) as **R** ∈ *ξ*. Edges on the dual network which connect cell centres are **T**_*j*_ *=* ∑_*i*_ *B*_*ij*_**R**_*i*_, *T*_*j*_ *=* |**T**_*j*_| and are orthogonal to **t**_*j*_ by construction.

**Fig. A1.**
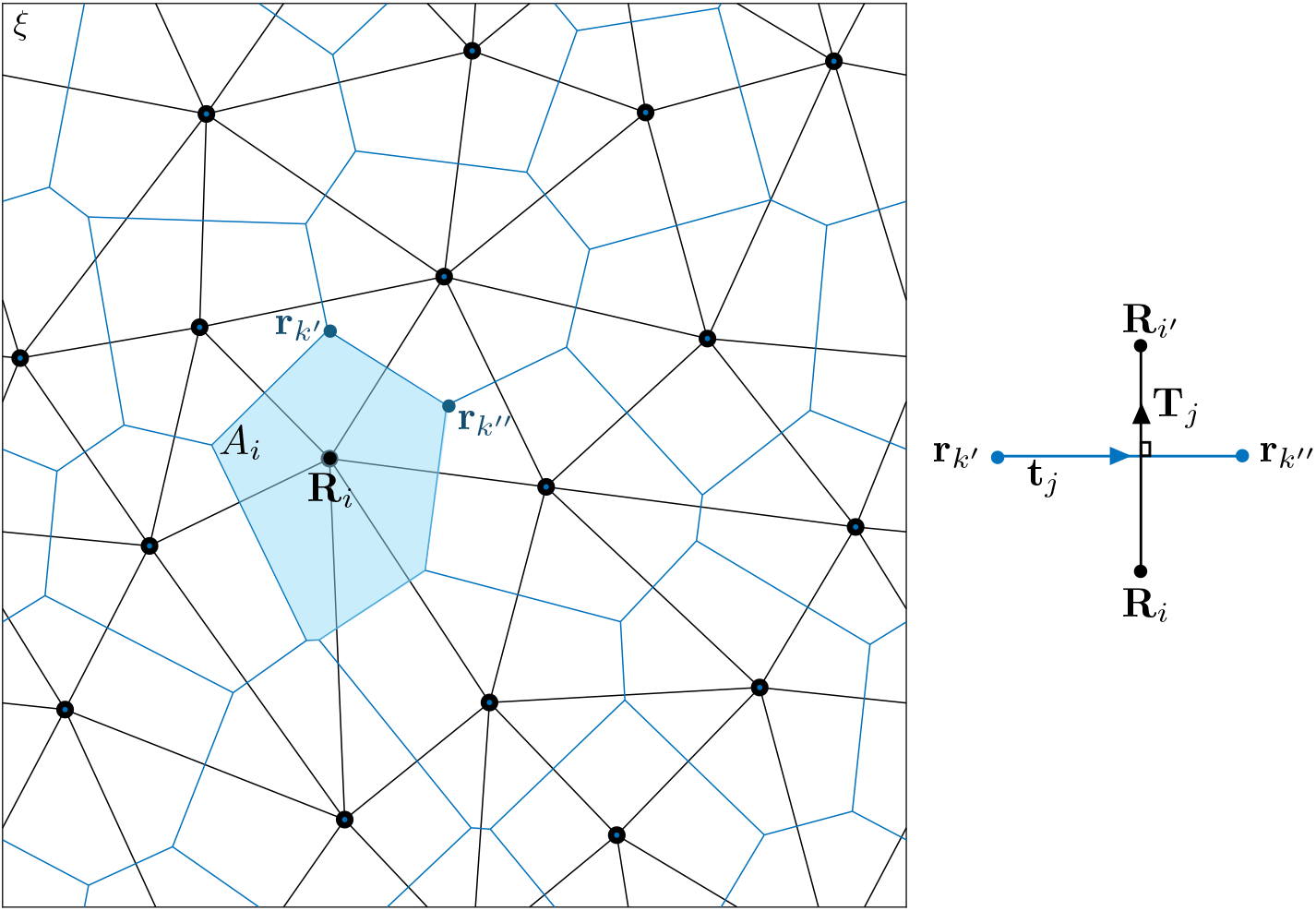
Depiction of a domain constructed as in A.2. We impose periodic boundaries.

We define the matrices H diag 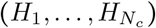, T diag 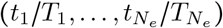, where *H*_*i*_ is the area of cell *i*, and the square matrix

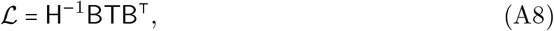

which is a Laplacian for scalar fields defined over cell centres [30], given the orthogonality of edges and links **t**_*j*_ ⋅ **T**_*j*_ *=* 0. We combine the GRN (2) in individual cells with Fickian diffusion between neighbouring cells using (A8) to obtain the reaction-diffusion equations (6). We nondimensionalise (6) by setting

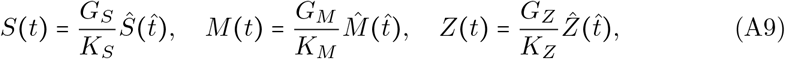

where 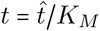. Dimensionless parameters are

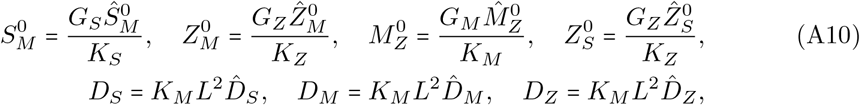

and 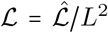, defining the length scale 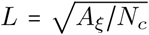, where *A*_*ξ*_ is the area of the domain *ξ* (in Figure A1).

### A.3 Steady uniform states

To determine the steady uniform states of (7) we solve the simultaneous equations (9). Solutions are *Ŝ* ∈ {1, *λ*_*ZS*_}, 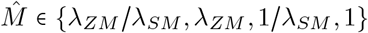, and 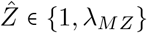, provided 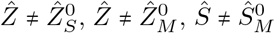, and 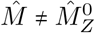. This gives rise to 16 possible steady uniform states. These states and the parameter restrictions that must be satisfied for feasibility are listed in Table A1. Further steady states will arise close to the discontinuities in the Heaviside functions under certain conditions. For example, if 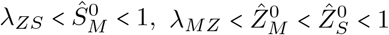 and 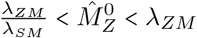, then the point 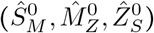 is a steady uniform state. For details see Tables A2 and A3. These states are listed at the end of Table A1 (SA1–SA3).

### A.4 Stability analysis

We determine the stability of steady uniform states to spatially-uniform perturbations using the Jacobian derived from (7)

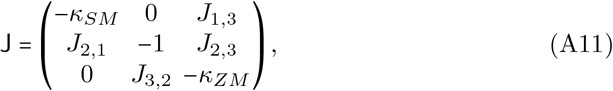

where

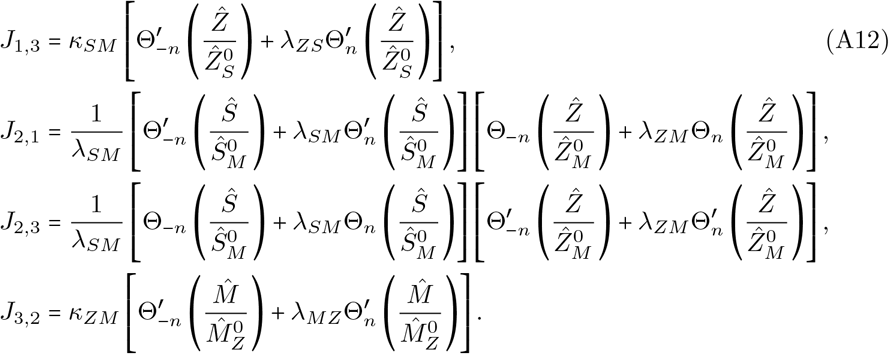

As *n* → ∞, 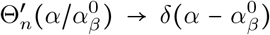, a Dirac delta function, so for *Ŝ* ∈ {1, *λ*_*ZS*_}, 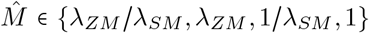 and 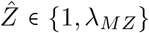 (values of *Ŝ*, 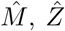 in states 1– 16 in Table 3), (A11) reduces to J = diag (−*κ*_*SM*_, −1, −*κ*_*ZM*_) and eigenvalues are the diagonal entries of J. They are negative and real, so states 1–16 are stable.

For the remaining states in Table 3, (A11) takes the form

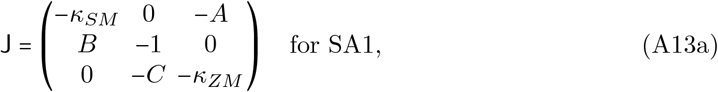

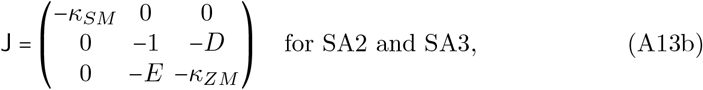

for some large values of *A, B, C, D, E >* 0 that grow linearly with *n*. The characteristic polynomials of (A13a) and (A13b) take the form

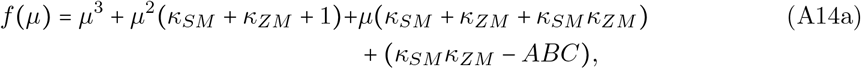

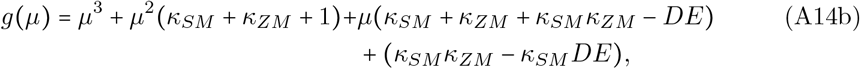

respectively. In the limit *n* → ∞,, *κ*_*SM*_ *κ*_*ZM*_ − *ABC <* 0, such that *f* (0) < 0. Together with lim_*µ*→∞_ *f* (*µ*) = ∞, there must be one positive real eigenvalue, *µ*_1_ > 0. Inspecting the signs of the sum and product of the remaining eigenvalues, we see that *µ*_2_ and *µ*_3_ either (i) form a complex conjugate pair with negative real part, or (ii) are negative and real. In either case, SA1 is a saddle point. Similar analysis of *g*(*µ*), with (*κ* +_*SM*_ *κ*_*ZM*_ + *κ*_*SM*_ *κ*_*ZM*_ − *DE*) < 0 and (*κ*_*SM*_ *κ*_*ZM*_ − *κ*_*SM*_ *DE*) < 0 as *n* → ∞, shows that SA2 and SA3 are also saddle points.

### A.5 Simulations

All 1D simulations are performed in Matlab using the variable-step variable-order solver ode15s. We will take *K*_*S*_ *= K*_*M*_ *= K*_*Z*_ and 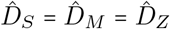. 2D simulations are performed in Julia, over the monolayer defined in A.2, using modified code from the Github repository by Revell *et al*. [48].

To estimate the speeds of travelling waves in Figures 5 and 6, we track a point along the wave in each variable. We choose the points 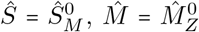 and 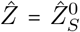. At each time point, we use linear interpolation to determine the horizontal position of the point of choice, and plot these positions against time. Since we have discretised space, we observe periodic bumps in space-time plots (a result of the linear interpolation). To approximate the speed of the travelling wave, we take the peaks of these bumps to calculate the wave speed, extracting data from a suitable time interval [*t*_0_, *t*_max_]. We overlay plots of the cell-level model to check that the shape does not change with time. We do this by using linear interpolation at *t = t*_0_ to estimate a value 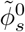 for which 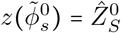. Along with the estimated wavespeed *c*, we overlay plots at a set of time points in the range *t*_*i*_ ∈ [*t*_0_, *t*_max_] by translating the wave along the horizontal axis by 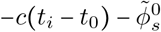. We translate by 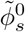 for reasons specified in Section 2.4.

## Appendix B Travelling-wave solutions

In this section we analyse travelling-wave solutions to the reaction-diffusion equations (7) in the 1D case, as described in Section 2.4 in the limit *n* → ∞.

**Table A1.** Steady states and the parameter restrictions which must be satisfied for each point to be consistent with the model. Labels are grouped by type (HD, TR, ML/NC, MS, HY, AL, M+, S+, SA), as defined in Table 2, and will be referenced in Section 3.1.

| Equilibrium Point $(\hat{S}, \hat{M}, \hat{Z})$ | Label | Parameter restrictions | | | |
| --- | --- | --- | --- | --- | --- |
| 1. $(\lambda_{ZS}, 1, \lambda_{MZ})$ | **HD1** | $\hat{S}_M^0 < \lambda_{ZS}$ | $\hat{M}_Z^0 < 1$ | $\hat{Z}_S^0 < \lambda_{MZ}$ | $\hat{Z}_M^0 > \lambda_{MZ}$ |
| 2. $(1, 1, \lambda_{MZ})$ | **HD2** | $\hat{S}_M^0 < 1$ | $\hat{M}_Z^0 < 1$ | $\hat{Z}_S^0 > \lambda_{MZ}$ | $\hat{Z}_M^0 > \lambda_{MZ}$ |
| 3. $(1, 1, 1)$ | **TR1** | $\hat{S}_M^0 < 1$ | $\hat{M}_Z^0 > 1$ | $\hat{Z}_S^0 > 1$ | $\hat{Z}_M^0 > 1$ |
| 4. $(1, \frac{1}{\lambda_{SM}}, 1)$ | **TR2** | $\hat{S}_M^0 > 1$ | $\hat{M}_Z^0 > \frac{1}{\lambda_{SM}}$ | $\hat{Z}_S^0 > 1$ | $\hat{Z}_M^0 > 1$ |
| 5. $(1, \lambda_{ZM}, 1)$ | **TR3** | $\hat{S}_M^0 < 1$ | $\hat{M}_Z^0 > \lambda_{ZM}$ | $\hat{Z}_S^0 > 1$ | $\hat{Z}_M^0 < 1$ |
| 6. $(1, \frac{1}{\lambda_{SM}}, \lambda_{MZ})$ | **ML1** | $\hat{S}_M^0 > 1$ | $\hat{M}_Z^0 < \frac{1}{\lambda_{SM}}$ | $\hat{Z}_S^0 > \lambda_{MZ}$ | $\hat{Z}_M^0 > \lambda_{MZ}$ |
| 7. $(1, \lambda_{ZM}, \lambda_{MZ})$ | **ML2** | $\hat{S}_M^0 < 1$ | $\hat{M}_Z^0 < \lambda_{ZM}$ | $\hat{Z}_S^0 > \lambda_{MZ}$ | $\hat{Z}_M^0 < \lambda_{MZ}$ |
| 8. $(1, \frac{\lambda_{ZM}}{\lambda_{SM}}, 1)$ | **NC** | $\hat{S}_M^0 > 1$ | $\hat{M}_Z^0 > \frac{\lambda_{ZM}}{\lambda_{SM}}$ | $\hat{Z}_S^0 < 1$ | $\hat{Z}_M^0 < 1$ |
| 9. $(\lambda_{ZS}, \frac{\lambda_{ZM}}{\lambda_{SM}}, 1)$ | **MS** | $\hat{S}_M^0 > \lambda_{ZS}$ | $\hat{M}_Z^0 > \frac{\lambda_{ZM}}{\lambda_{SM}}$ | $\hat{Z}_S^0 < 1$ | $\hat{Z}_M^0 < 1$ |
| 10. $(\lambda_{ZS}, 1, 1)$ | **HY1** | $\hat{S}_M^0 < \lambda_{ZS}$ | $\hat{M}_Z^0 > 1$ | $\hat{Z}_S^0 < 1$ | $\hat{Z}_M^0 > 1$ |
| 11. $(\lambda_{ZS}, \lambda_{ZM}, 1)$ | **HY2** | $\hat{S}_M^0 < \lambda_{ZS}$ | $\hat{M}_Z^0 > \lambda_{ZM}$ | $\hat{Z}_S^0 < 1$ | $\hat{Z}_M^0 < 1$ |
| 12. $(\lambda_{ZS}, \frac{1}{\lambda_{SM}}, 1)$ | **HY3** | $\hat{S}_M^0 > \lambda_{ZS}$ | $\hat{M}_Z^0 > \frac{1}{\lambda_{SM}}$ | $\hat{Z}_S^0 < 1$ | $\hat{Z}_M^0 > 1$ |
| 13. $(\lambda_{ZS}, \frac{\lambda_{ZM}}{\lambda_{SM}}, \lambda_{MZ})$ | **AL** | $\hat{S}_M^0 > \lambda_{ZS}$ | $\hat{M}_Z^0 < \frac{\lambda_{ZM}}{\lambda_{SM}}$ | $\hat{Z}_S^0 < \lambda_{MZ}$ | $\hat{Z}_M^0 < \lambda_{MZ}$ |
| 14. $(\lambda_{ZS}, \frac{1}{\lambda_{SM}}, \lambda_{MZ})$ | **M+1** | $\hat{S}_M^0 > \lambda_{ZS}$ | $\hat{M}_Z^0 < \frac{1}{\lambda_{SM}}$ | $\hat{Z}_S^0 < \lambda_{MZ}$ | $\hat{Z}_M^0 > \lambda_{MZ}$ |
| 15. $(\lambda_{ZS}, \lambda_{ZM}, \lambda_{MZ})$ | **M+2** | $\hat{S}_M^0 < \lambda_{ZS}$ | $\hat{M}_Z^0 < \lambda_{ZM}$ | $\hat{Z}_S^0 < \lambda_{MZ}$ | $\hat{Z}_M^0 < \lambda_{MZ}$ |
| 16. $(1, \frac{\lambda_{ZM}}{\lambda_{SM}}, \lambda_{MZ})$ | **S+** | $\hat{S}_M^0 > 1$ | $\hat{M}_Z^0 < \frac{\lambda_{ZM}}{\lambda_{SM}}$ | $\hat{Z}_S^0 > \lambda_{MZ}$ | $\hat{Z}_M^0 < \lambda_{MZ}$ |
| 17. $(\hat{S}_M^0, \hat{M}_Z^0, \hat{Z}_S^0)$ | **SA1** | | | | |
| 18. $(\lambda_{ZS}, \hat{M}_Z^0, \hat{Z}_M^0)$ | **SA2** | See Tables A2 and A3. | | | |
| 19. $(1, \hat{M}_Z^0, \hat{Z}_M^0)$ | **SA3** | | | | |

### B.1 ODE system

We introduce the variables *v*_*s*_ (*ϕ*), *v*_*m*_ (*ϕ*), *v*_*z*_ (*ϕ*) to rewrite (11) as a 6-dimensional system of first-order ODEs:

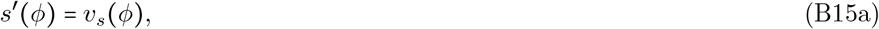

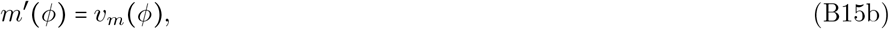

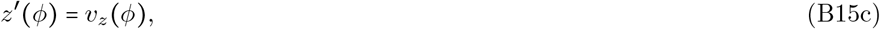

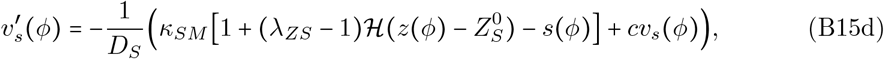

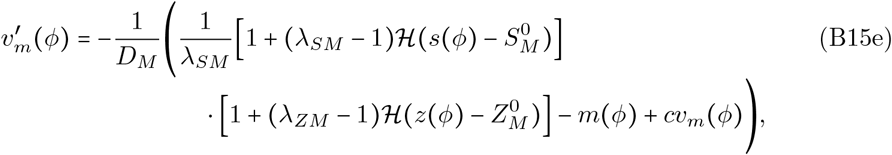

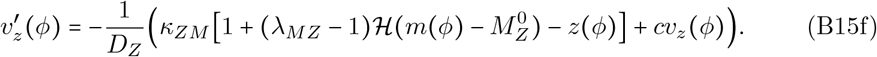

**Table A2.**
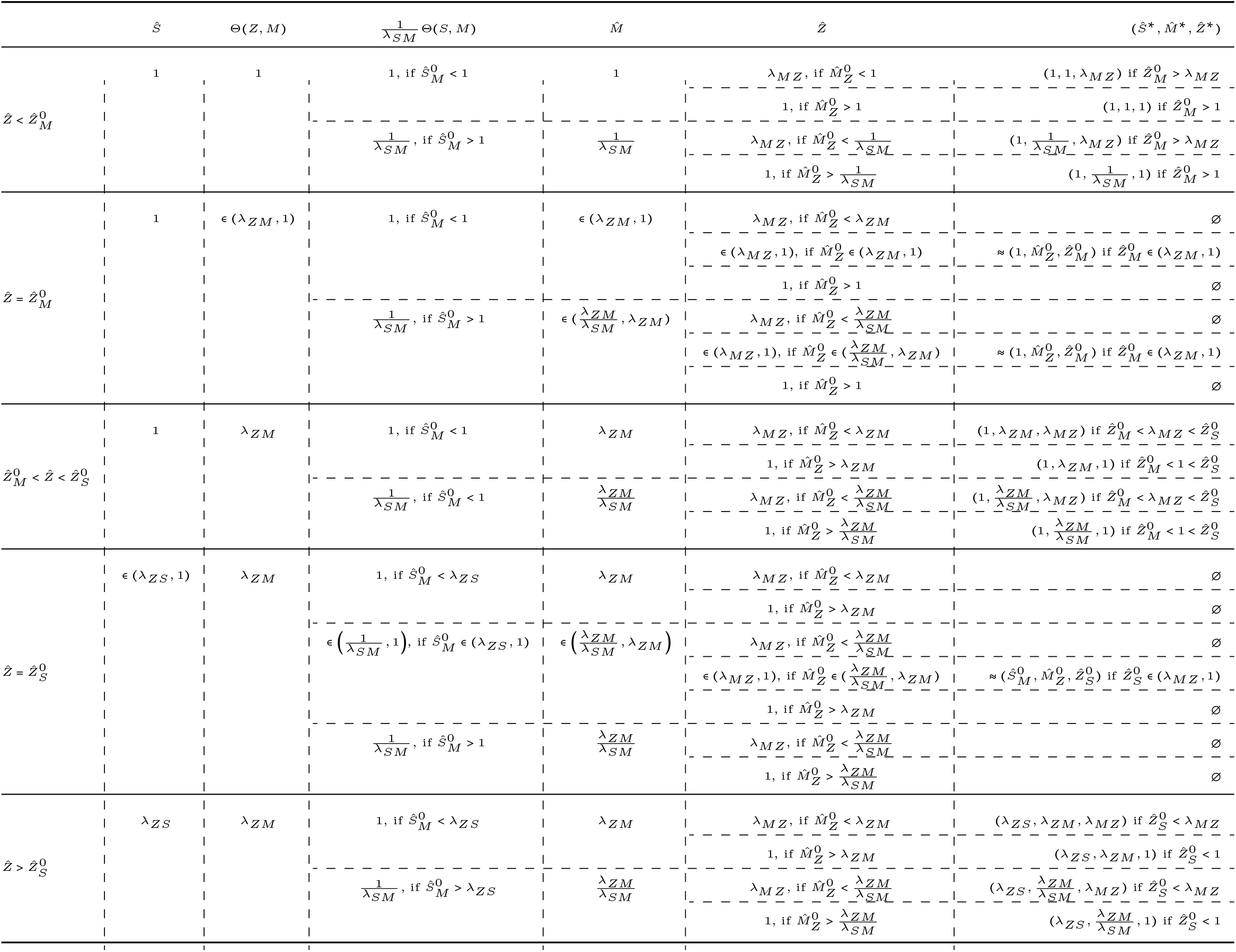
Table of solutions to the simultaneous equations (9) as 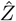 varies, for 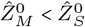.

**Table A3.**
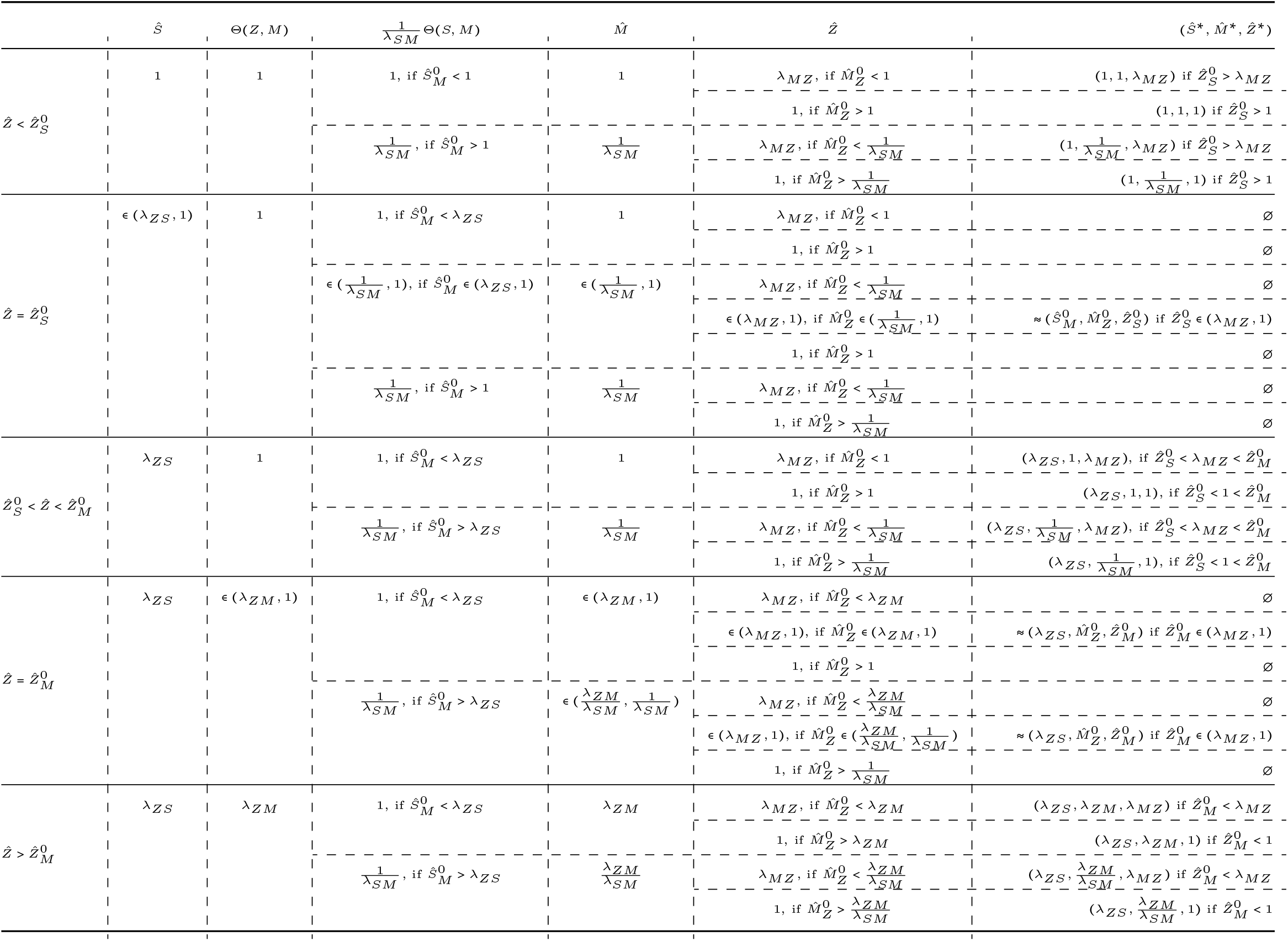
Table of solutions to the simultaneous equations (9) as 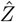 varies, for 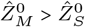.

### B.2 Exponential coefficients

The piecewise-exponential travelling-wave solution (18) is constructed as described in Section 2.4 and using the information in Table B4 to reduce the number of unknown coefficients, determined as follows. Taking 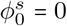 in the travelling-wave solution (18) and using boundary conditions uncovers the following set of 17 simultaneous equations with 17 unknowns (namely 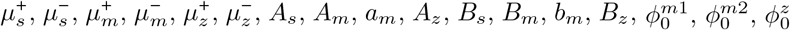):

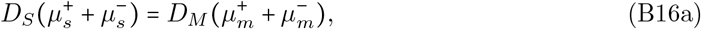

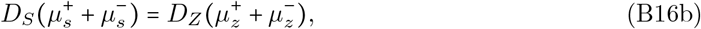

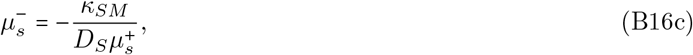

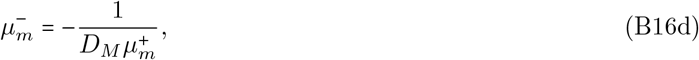

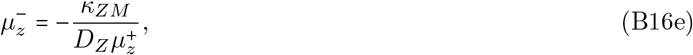

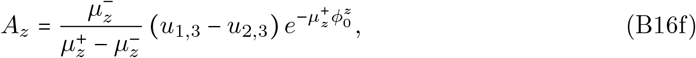

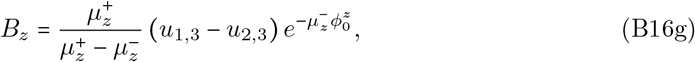

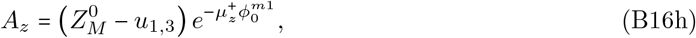

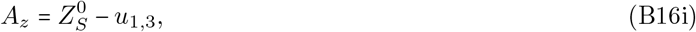

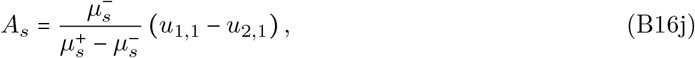

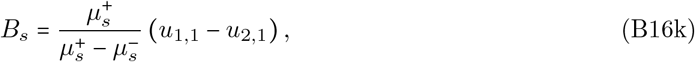

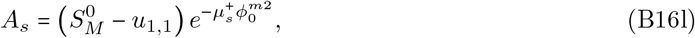

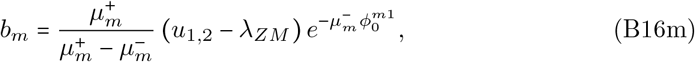

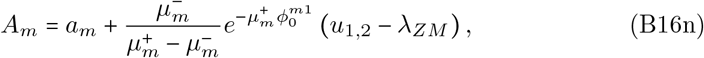

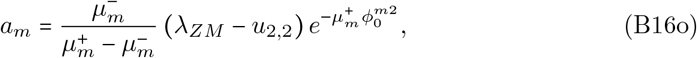

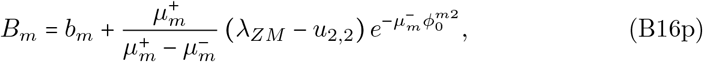

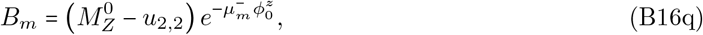

where we have taken 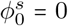 in (B16i), (B16j) and (B16k). (B16a)–(B16e) arise from the dispersion relations (19); (B16f), (B16g) from patching *z*(*ϕ*) and *z ′* (*ϕ*) at 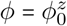; (B16h) from the condition 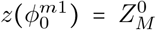; (B16i) from the condition 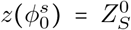; (B16j),(B16k) from patching *s*(*ϕ*) and *s*^′^(*ϕ*) at 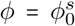; (B16l) from the condition 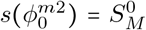; (B16m), (B16n) from patching *m*(*ϕ*) and *m*^′^(*ϕ*) at 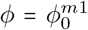; (B16o), (B16p) from patching *m*(*ϕ*) and *m′* (*ϕ*) at 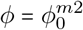; (B16q) from the condition 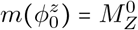.

**Table B4.**
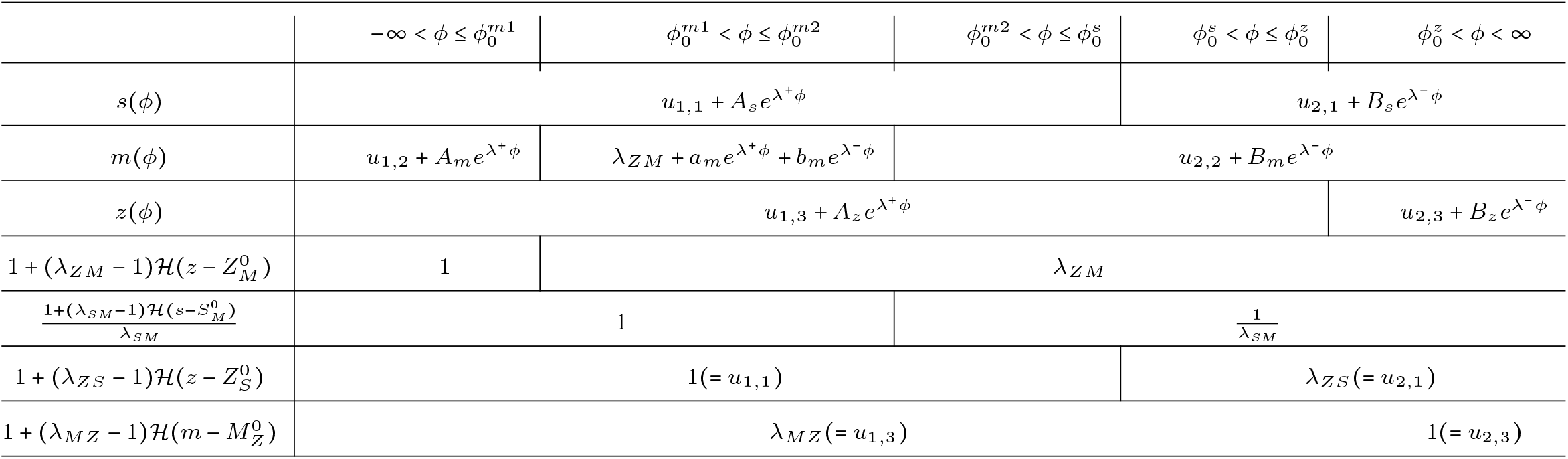
Table showing which components of the exponential solution are affected as each step function changes. We divide the *ϕ*-axis into five regions, separated by the switching thresholds 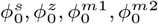. This is used to construct the piecewise exponential solution in (18).

### B.3 Spatiotemporal stability threshold outside of Ω

Here we give an overview of how the spatiotemporal stability thresholds in Figure 3(d) are derived. Section 2.4 and Appendix B give the derivation for parameters in Ω (21), and thus for 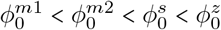 (17). The order and existence of these *ϕ* will affect the following equations in (B16):

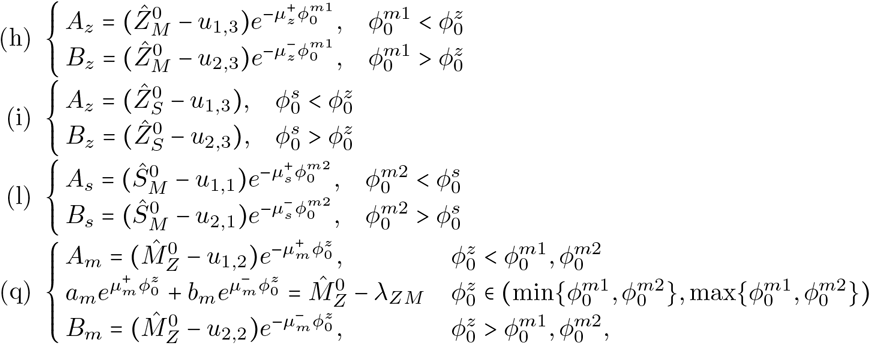

and 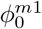 and 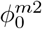 will be swapped in (m),(n) and (o),(p), respectively, if 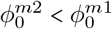. We determine the order of these *ϕ* by tracking threshold values on the curve as described in Section A.5 for desired parameters. Furthermore, if any threshold value does not lie between the two equilibria values in a given parameter region, the associated *ϕ* will be removed from the system of equations. For example, if 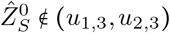, then 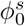 is not defined and we eliminate equations (B16)(i),(j),(k). The resulting system of equations is solved for the stationary case *c =* 0 and the spatiotemporal stability threshold is derived.

## Appendix C Robustness for *n*< ∞

### C.1 Bifurcation diagrams

We consider the robustness of the results in Section 3.1 for *n* < ∞. Figure 3 reveals saddle-node bifurcations as parameters 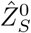 or 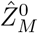 cross certain thresholds, while 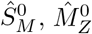 are kept constant. Figure 3 shows a clearly defined region within which the system is bistable. We consider how the shape of this region changes for finite values of *n* in Figure C2. Using the Julia package *BifurcationKit*.*jl* [49] we perform continuation in 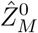 and 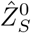 from initial guesses based on our approximation *n* → ∞, plotting the contour of the bifurcation boundary. We see in Figure C2 that for *n* ≤ 6 the results are sensitive to the value of *n*: the bistable region disappears for *n* ≤ 2 and we see some cusping behaviour.

At *n* 5, we observe a butterfly catastrophe, a depiction and explanation of which can be found in [34]. There is a small region of parameter space within which the system (7) has 5 steady uniform states. In Figure C2(b) we consider fractional values of *n* between *n =* 4 and *n =* 5 to see how the bifurcation boundaries evolve to this structure, which is reminiscent of the structures seen in [34, Figure 4.11, p. 55]. In Figure C2(c) we plot the evolution of the bifurcation boundary for larger integer values of *n*, and observe that the structure quickly approaches the dashed line (representing the limit as *n* → ∞, from Figure 3(b)).

**Fig. C2.**
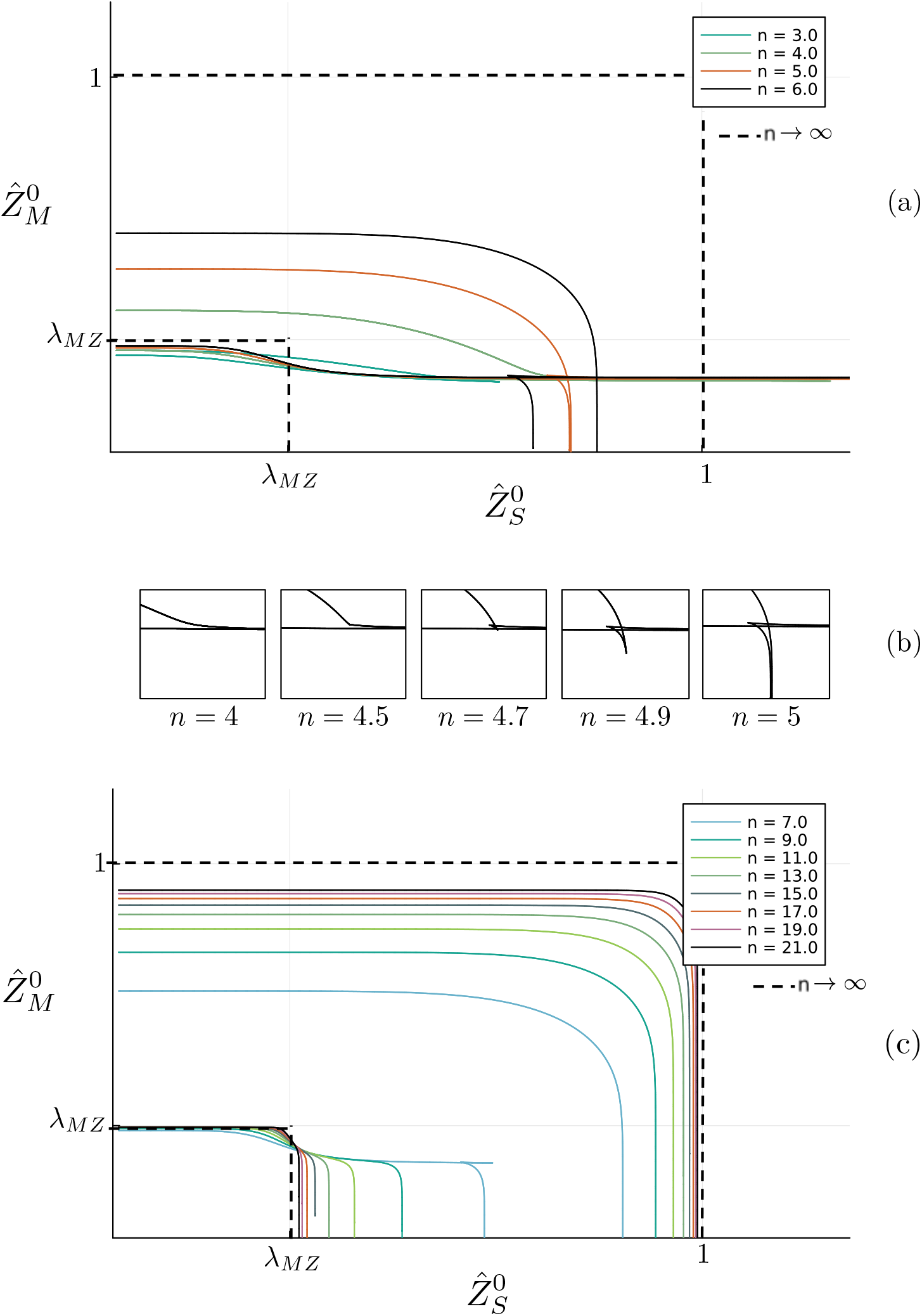
Bifurcation boundary contours from Figure 3(b) for finite values of n. Panel (a) shows contours for integer values *n ≤* 6, with the boundary disappearing for *n ≤* 2 and cusping behaviour is observed. (b) shows a zoom-in of the cusp formation between *n* = 4 and *n* = 5, with evidence of a butterfly catastrophe. (c) shows the boundary for integer values 7 ≤ *n* ≤ 21, and we see the boundaries quickly approaching the dashed line, the limit *n* → ∞.

## Notes

https://github.com/charlietaylorbarca1/Bifurcation_Plots

https://github.com/charlietaylorbarca1/GRN_Simulations

https://github.com/charlietaylorbarca1/Monolayer_Construction

https://github.com/charlietaylorbarca1/DemoForCharlie

### Competing Interest Statement

The authors have declared no competing interest.

### Summary of Updates

Section 1 updated to clarify modelling assumptions, and expand the mathematical literature review; Section 2 updated to clarify boundary conditions and assumptions In travelling wave analysis; section 3 updated to consider a variety of diffusion coefficients; discussion updated to comment on further on future research directions and conclusions from the model; typographical and reference edits corrected throughout.

## References

[1] Wolf, Y., Bartok, O., Patkar, S., Eli, G.B., Cohen, S., Litchfield, K., Levy, R., Jiménez-Sánchez, A., Trabish, S., Lee, J.S., et al.: UVB-induced tumor heterogeneity diminishes immune response in melanoma. Cell 179, 219–235 (2019)

[2] Carreira, S., Goodall, J., Denat, L., Rodriguez, M., Nuciforo, P., Hoek, K.S., Testori, A., Larue, L., Goding, C.R.: MITF regulation of Dia1 controls melanoma proliferation and invasiveness. Genes Dev. 20, 3426–3439 (2006)

[3] Hoek, K.S., Goding, C.R.: Cancer stem cells versus phenotype-switching in melanoma. Pigment Cell Melanoma Res. 23, 746–759 (2010)

[4] Rambow, F., Marine, J.-C., Goding, C.R.: Melanoma plasticity and phenotypic diversity: therapeutic barriers and opportunities. Genes Dev. 33, 1295–1318 (2019)

[5] Goding, C.R., Arnheiter, H.: MITF—the first 25 years. Genes Dev. 33, 983–1007 (2019)

[6] Karras, P., Bordeu, I., Pozniak, J., Nowosad, A., Pazzi, C., Van Raemdonck, N., Landeloos, E., Van Herck, Y., Pedri, D., Bervoets, G., et al.: A cellular hierarchy in melanoma uncouples growth and metastasis. Nature 610, 190–198 (2022)

[7] Tsoi, J., Robert, L., Paraiso, K., Galvan, C., Sheu, K.M., Lay, J., Wong, D.J., Atefi, M., Shirazi, R., Wang, X., et al.: Multi-stage differentiation defines melanoma subtypes with differential vulnerability to drug-induced iron-dependent oxidative stress. Cancer Cell 33, 890–904 (2018)

[8] Taylor, K.L., Lister, J.A., Zeng, Z., Ishizaki, H., Anderson, C., Kelsh, R.N., Jackson, I.J., Patton, E.E.: Differentiated melanocyte cell division occurs in vivo and is promoted by mutations in MITF. Development 138, 3579–3589 (2011)

[9] Durand, S., Tang, Y., Pommier, R.M., Benboubker, V., Grimont, M., Boivin, F., Barbollat-Boutrand, L., Cumunel, E., Dupeuble, F., Eberhardt, A., et al.: ZEB1 controls a lineage-specific transcriptional program essential for melanoma cell state transitions. Oncogene 43, 1489–1505 (2024)

[10] Plaschka, M., Benboubker, V., Grimont, M., Berthet, J., Tonon, L., Lopez, J., Le-Bouar, M., Balme, B., Tondeur, G., Fouchardiére, A., et al.: ZEB1 transcription factor promotes immune escape in melanoma. J. Immunother. Cancer 10 (2022)

[11] Denecker, G., Vandamme, N., Akay, Ö., Koludrovic, D., Taminau, J., Lemeire, K., Gheldof, A., De Craene, B., Van Gele, M., Brochez, L., et al.: Identification of a ZEB2-MITF-ZEB1 transcriptional network that controls melanogenesis and melanoma progression. Cell Death Differ. 21, 1250–1261 (2014)

[12] Chen, Y., Lu, X., Montoya-Durango, D.E., Liu, Y.-H., Dean, K.C., Darling, D.S., Kaplan, H.J., Dean, D.C., Gao, L., Liu, Y.: ZEB1 regulates multiple oncogenic components involved in uveal melanoma progression. Sci. Rep. 7, 45 (2017)

[13] Bondurand, N., Pingault, V., Goerich, D.E., Lemort, N., Sock, E., Caignec, C.L., Wegner, M., Goossens, M.: Interaction among SOX10, PAX3 and MITF, three genes altered in Waardenburg syndrome. Hum. Mol. Genet. 9, 1907–1917 (2000)

[14] Dilshat, R., Fock, V., Kenny, C., Gerritsen, I., Lasseur, R.M.J., Travnickova, J., Eichhoff, O.M., Cerny, P., Möller, K., Sigurbjörnsdóttir, S., et al.: MITF reprograms the extracellular matrix and focal adhesion in melanoma. eLife 10, 63093 (2021)

[15] Liu, Y., Ye, F., Li, Q., Tamiya, S., Darling, D.S., Kaplan, H.J., Dean, D.C.: Zeb1 represses MITF and regulates pigment synthesis, cell proliferation, and epithelial morphology. Invest. Ophthal. Vis. Sci. 50, 5080–5088 (2009)

[16] Gopalan, V., Wong, C.W., Leshem, R., Owen, L., Vallius, T., Shi, Y., Jiang, Y., Pérez-Guijarro, E., Wu, E., Chin, S., et al.: Transitory Schwann Cell Precursor and hybrid states underpin melanoma therapy resistance and metastasis. bioRxiv (2025) 10.1101/2022.10.14.512297

[17] Pham, F., Dufeu, M., Benboubker, V., Grimont, M., Lhorisson, A., Berthet, J., Donzel, M., Schneider, R., Tonon, L., Doffin, A.-C., et al.: Spatial tumour-immune ecosystems shape the efficacy of anti-PD1 immunotherapy in primary cutaneous melanoma. bioRxiv (2025)

[18] Vallius, T., Shi, Y., Novikov, E., Pant, S.M., Pelletier, R., Chen, Y.-A., Tefft, J.B., Johnson, A.N., Maliga, Z., Wan, G., et al.: Spatial determinants of tumor cell dedifferentiation and plasticity in primary cutaneous melanoma. bioRxiv (2025)

[19] Subhadarshini, S., Sahoo, S., Debnath, S., Somarelli, J.A., Jolly, M.K.: Dynamical modeling of proliferative-invasive plasticity and IFNγ signaling in melanoma reveals mechanisms of PD-L1 expression heterogeneity. J. Immunother. Cancer 11(9) (2023)

[20] Casey, R., Jong, H.D., Gouzé, J.L.: Piecewise-linear models of genetic regulatory networks: Equilibria and their stability. J. Math. Biol. 52, 27–56 (2006)

[21] Plahte, E., Kjøglum, S.: Analysis and generic properties of gene regulatory networks with graded response functions. Physica D: Nonlinear Phenomena 201, 150–176 (2005)

[22] Santillán, M.: On the use of the Hill functions in mathematical models of gene regulatory networks. Math. Model. Nat. Phenom. 3(2), 85–97 (2008)

[23] Wolpert, L.: One hundred years of positional information. Trends in Genetics 12, 359–364 (1996)

[24] Barbier, I., Perez-Carrasco, R., Schaerli, Y.: Controlling spatiotemporal pattern formation in a concentration gradient with a synthetic toggle switch. Mol. Sys. Biol. 16, 199361 (2020)

[25] Roy, U., Singh, D., Vincent, N., Haritas, C.K., Jolly, M.K.: Spatiotemporal patterning enabled by gene regulatory networks. ACS omega 8, 3725 (2023)

[26] Diambra, L., Senthivel, V.R., Menendez, D.B., Isalan, M.: Cooperativity to increase turing pattern space for synthetic biology. ACS Synthetic Biology 4, 177–186 (2015)

[27] Murray, J.D.: Mathematical Biology : An Introduction. Springer, New York (2002)

[28] Dang, Y., Grundel, D.A.J., Youk, H.: Cellular dialogues: Cell-cell communication through diffusible molecules yields dynamic spatial patterns. Cell Systems 10, 82–987 (2020)

[29] Gelens, L., Anderson, G.A., Ferrell Jr, J.E.: Spatial trigger waves: positive feedback gets you a long way. Mol. Biol. Cell 25, 3486–3493 (2014)

[30] Jensen, O.E., Revell, C.K.: Couple stresses and discrete potentials in the vertex model of cellular monolayers. Biomech. Model. Mechanobiol. 22, 1465–1486 (2023)

[31] Rosen, G.: On the Fisher and the cubic-polynomial equations for the propagation of species properties. Bull. Math. Biol. 42, 95–106 (1980)

[32] Rambow, F., Rogiers, A., Marin-Bejar, O., Aibar, S., Femel, J., Dewaele, M., Karras, P., Brown, D., Chang, Y.H., Debiec-Rychter, M., et al.: Toward minimal residual disease-directed therapy in melanoma. Cell 174, 843–855 (2018)

[33] Pozniak, J., Pedri, D., Landeloos, E., Van Herck, Y., Antoranz, A., Vanwynsberghe, L., Nowosad, A., Roda, N., Makhzami, S., Bervoets, G., et al.: A TCF4-dependent gene regulatory network confers resistance to immunotherapy in melanoma. Cell 187, 166–183 (2024)

[34] Saunders, P.T.: An Introduction to Catastrophe Theory. Cambridge University Press, Cambridge (1980)

[35] Marie, K.L., Sassano, A., Yang, H.H., Michalowski, A.M., Michael, H.T., Guo, T., Tsai, Y.C., Weissman, A.M., Lee, M.P., Jenkins, L.M., et al.: Melanoblast transcriptome analysis reveals pathways promoting melanoma metastasis. Nature Communications 11(1), 333 (2020)

[36] Oyama, T., Ogawa, H., Shirai, Y., Abe, H., Kamiya, T., Abe, T., Tanuma, S.-i.: Hinokitiol-induced decreases of tyrosinase and microphthalmia-associated transcription factor are mediated by the endoplasmic reticulum-associated degradation pathway in human melanoma cells. Biochimie 192, 13–21 (2022)

[37] Munteanu, C.V., Chirit oiu, G.N., Chirit oiu, M., Ghenea, S., Petrescu, A.-J., Petrescu, Ş.M.: Affinity proteomics and deglycoproteomics uncover novel edem2 endogenous substrates and an integrative erad network. Molecular & Cellular Proteomics 20, 100125 (2021)

[38] Hansen, A.S., O’Shea, E.K.: Cis determinants of promoter threshold and activation timescale. Cell Rep. 12, 1226–1233 (2015)

[39] Liguori, L., Luciano, A., Pagliara, V., Polcaro, G., De Feo, R., Viggiano, A., Salomone, F., Avanzo, A., Vitale, F., D’ambrosio, S., et al.: Epigenetic modifiers to enhance the efficacy of immune checkpoint inhibitors for the treatment of melanoma. Transl. Oncol. 59, 102452 (2025)

[40] Hunter, M.V., Joshi, E., Bowker, S., Montal, E., Ma, Y., Kim, Y.H., Yang, Z., Tuffery, L., Li, Z., Rosiek, E., et al.: Mechanical confinement governs phenotypic plasticity in melanoma. Nature 647, 517–527 (2025)

[41] Pisco, A.O., Huang, S.: Non-genetic cancer cell plasticity and therapy-induced stemness in tumour relapse:’what does not kill me strengthens me’. Brit. J. Cancer 112, 1725–1732 (2015)

[42] Eom, J., Park, S.M., Feisst, V., Chen, C.J.J., Mathy, J.E., McIntosh, J.D., Angel, C.E., Bartlett, A., Martin, R., Mathy, J.A., Cebon, J.S., Black, M.A., Brooks, A.E.S., Dunbar, P.R.: Distinctive subpopulations of stromal cells are present in human lymph nodes infiltrated with melanoma. Cancer Immunol. Res. 8, 990– 1003 (2020)

[43] Huang, B., Lu, M., Jia, D., Ben-Jacob, E., Levine, H., Onuchic, J.N.: Interrogating the topological robustness of gene regulatory circuits by randomization. PLoS Comp. Biol. 13(3), 1005456 (2017)

[44] Harris, R.E., Pargett, M., Sutcliffe, C., Umulis, D., Ashe, H.L.: Brat promotes stem cell differentiation via control of a bistable switch that restricts BMP signaling. Dev. Cell 20, 72–83 (2011)

[45] Olimpio, E.P., Dang, Y., Youk, H.: Statistical dynamics of spatial-order formation by communicating cells. Iscience 2, 27–40 (2018)

[46] Coppey, M., Berezhkovskii, A.M., Sealfon, S.C., Shvartsman, S.Y.: Time and length scales of autocrine signals in three dimensions. Biophys. J. 93(6), 1917– 1922 (2007)

[47] Grady, L.J., Polimeni, J.R.: Discrete Calculus: Applied Analysis on Graphs for Computational Science vol. 3. Springer, London (2010)

[48] Revell, C.K., Jensen, O.E.: VertexModel.jl (2022). https://github.com/chris-revell/VertexModel

[49] Veltz, R.: BifurcationKit.jl (2020). https://hal.archives-ouvertes.fr/hal-02902346

